# Human cerebral cortex networks use expanding and contracting state dynamic to shape cortical functions

**DOI:** 10.1101/2020.09.24.309468

**Authors:** Alex Willumsen, Jens Midtgaard, Bo Jespersen, Christoffer K.K. Hansen, Salina N. Lam, Sabine Hansen, Ron Kupers, Martin E. Fabricius, Minna Litman, Lars Pinborg, José D. Tascón-Vidarte, Anne Sabers, Per E. Roland

## Abstract

We lack viable explanations of how brain functions emerge from collective activities of neurons in networks. We recorded field potentials from many local networks in the human cerebral cortex during a wide variety of brain functions. The network dynamics showed that each local cortical network produced fluctuating attractor states. The state trajectories continuously stretched and contracted during all brain functions, leaving no stable patterns. Different local networks all produced this dynamic, despite different architectures. Single trial stimuli and tasks modified the stretching and contractions. These modified fluctuations cross-correlated among particular networks during specific brain functions. Spontaneous states, rest, sensory, motor and cognitive states all emerged from this dynamic. Its mathematical structure provides a general explanation of cortical dynamics that can be tested experimentally. This universal dynamic is a simple functional organizing principle for brain functions at the mm^3^ scale that is distinct from existing frameworks.

---

Cognition, behavior and perception emerge from collective action of neurons in local networks and across networks in the cerebral cortex and other parts of the brain. The cortical operations supporting these larger scale collective actions are not known (Breakspear, 2017; Buzsaki, 2019; Ju and Bassett 2020, McCormick et al., 2020; Singer et al., 2019).

The purpose of this study is to reveal the dynamics of cortical operations at a larger scale. Local cortical networks contribute to cortical operations by changing their cortical states. The dynamics of a local network explains the temporal evolution of the cortical states it generates (Babloyantz and Destexhe, 1986; Breakspear, 2017; Ju and Bassett, 2020). Examining the dynamics exhaustively from many local networks will show how their changes of cortical states support cortical operations leading to cognition, behavior and perception. Such an analysis may also reveal if there are general mechanisms underlying cortical operations.

Substantial parts of cortical operations are carried out by postsynaptic operations of neurons in local cortical networks (Rheinberger and Jasper 1937; Iacaruso et al. 2018; McCormick et al., 2020). The postsynaptic membrane currents in a local cortical network collectively generate a field potential. The field potential thus contains information of collective postsynaptic operations at a larger mm^3^ spatial scale (Einevoll et al., 2014; Pesaran et al., 2018). Examining the field potential dynamics captures the non-linear behavior of these postsynaptic operations during a brain function, also in single trials. This is not possible with the traditional analysis, in which field potentials are separated into different frequency bandwidths under the assumption of stationarity (Babloyantz and Destexhe, 1986; Breakspear, 2017; Rabinovich et al., 2012; Roschke and Basar,1990; Stam, 1996).

Different types dynamical models have been proposed to explain the dynamics in field potentials during one or a few conditions in selected parts of the cortex, but with no consensus (Tognioli and Kelso 2011; Knierim and Zhang, 2012; Rabinovich et al., 2012; Priesemann et al., 2013; Wimmer and Nykamp, 2014; Breakspear, 2017; Hennequin et al., 2018; Ju and Bassett, 2020). This could mean that different types of dynamics are reserved for different cortical functions. It could also mean that cortical networks, having different architectures, produce different dynamics. Models must be supported by general experimental evidence and they must account for the ever irregularly self-sustained fluctuations of postsynaptic membrane currents and field potentials, continuously changing states everywhere in the cortex (Rheinberger and Jasper,1937; Einevoll et al., 2014; Pesaran et al., 2018; McCormick et al., 2020).

The field potential is multi-dimensional, because it is created by many currents in the local network. Each measurement of the field potential corresponds to one point in a multidimensional space (scaled in μV). This point is a cortical state of the local network. Here we developed a method for exhaustively deriving the dynamics directly from the empirical field potential without unrealistic assumptions. This enabled us to reveal the dynamics used by many local cortical networks during spontaneous activity and while patients performed many sensory, motor and cognitive tasks.

Despite different architectures, all probed local networks produced successions of local cortical states irregularly stretching and contracting their trajectories. The trajectories were located to an attractor that consequently expanded and contracted unceasingly. This unique combination of attractor stability with vigorous fluctuating expanding and contracting states drove cortical operations locally and between networks, also in single trials. Moreover, despite the large diversity of brain functions tested, the underlying operations all evolved from variations of these cortical state fluctuations. Given the diversity of local cortical networks and cortical functions, these conclusions may seem counterintuitive. However, our results are solid and reproducible. They explain how changes in local cortical states can be shared by many networks. They provide first empirical evidence for a general dynamic governing local and larger scale cortical operations at functionally relevant time scales, since the proposal of synchrony (Gray and Singer, 1989). Having only one type of dynamics opens for more specific hypotheses and studies of cortical operations, which can be directly tested experimentally in animals and also with MEG and EEG.

## RESULTS

We recorded field potentials from multiple sites in the cerebral cortex of 13 patients with medical intractable epilepsy (**Methods**). In a local cortical network, the field potential is dominated by transmembrane currents from more than 10^5^ dendrites and their subsequent extracellular currents (Einevoll et al., 2014; Pesaran et al., 2018; **Methods**). Each field potential time-series reflects the underlying dynamics of the local network in the immediate neighborhood of the electrode lead at which it is recorded. Each analyzed time series was 60s long (**Methods; Figure S1**). All epileptic activity was removed and the analysis was restricted to those leads and time series that remained without any abnormality throughout the experiments. This left 3476 60s files of normal field potential recordings, distributed over 290 electrode leads under 40 different conditions. These conditions comprised visual, auditory, gustatory, olfactory and somatosensory stimulations, as well as tests with voluntary movements in which the dynamics of the brain operations was driven partly by external cues or stimuli. We also included memory retrieval and mental navigation tasks that did not involve external stimuli and overt behavior (**Methods**).

### Dimensionality of field potential dynamics

The dimensionality is the number of dimensions we need to get an exhaustive account of the information about cortical states in the field potential (Roschke and Basar, 1990; Stam 1996). Although the field potential is multidimensional, its true dimensionality is unknown. We therefore need first to determine the number of dimensions needed to create the multidimensional space, in which we can study how the cortical states produced by each local network evolve. First, we embedded the field potential (Takens, 1981) (**Figures 1 and S1; Methods**). Then, we estimated its correlation dimensionality. To find the dynamics driving the evolution of the multi-dimensional field potential in one local network implies to demonstrate how the local cortical states produced by the local network evolve in a multidimensional state space of sufficient dimensions. A local cortical state is one point in this multidimensional state space. In the state spaces in this study, one point represents the state of the local network over 1.95 ms (**Figure 1**). Despite the intrinsic high dimensionality of the sources of the field potential, the estimated dimensionality was always comparatively low, ranging from 3 to 9 (**Figures 2 and S2)**.

**Figure 1.**
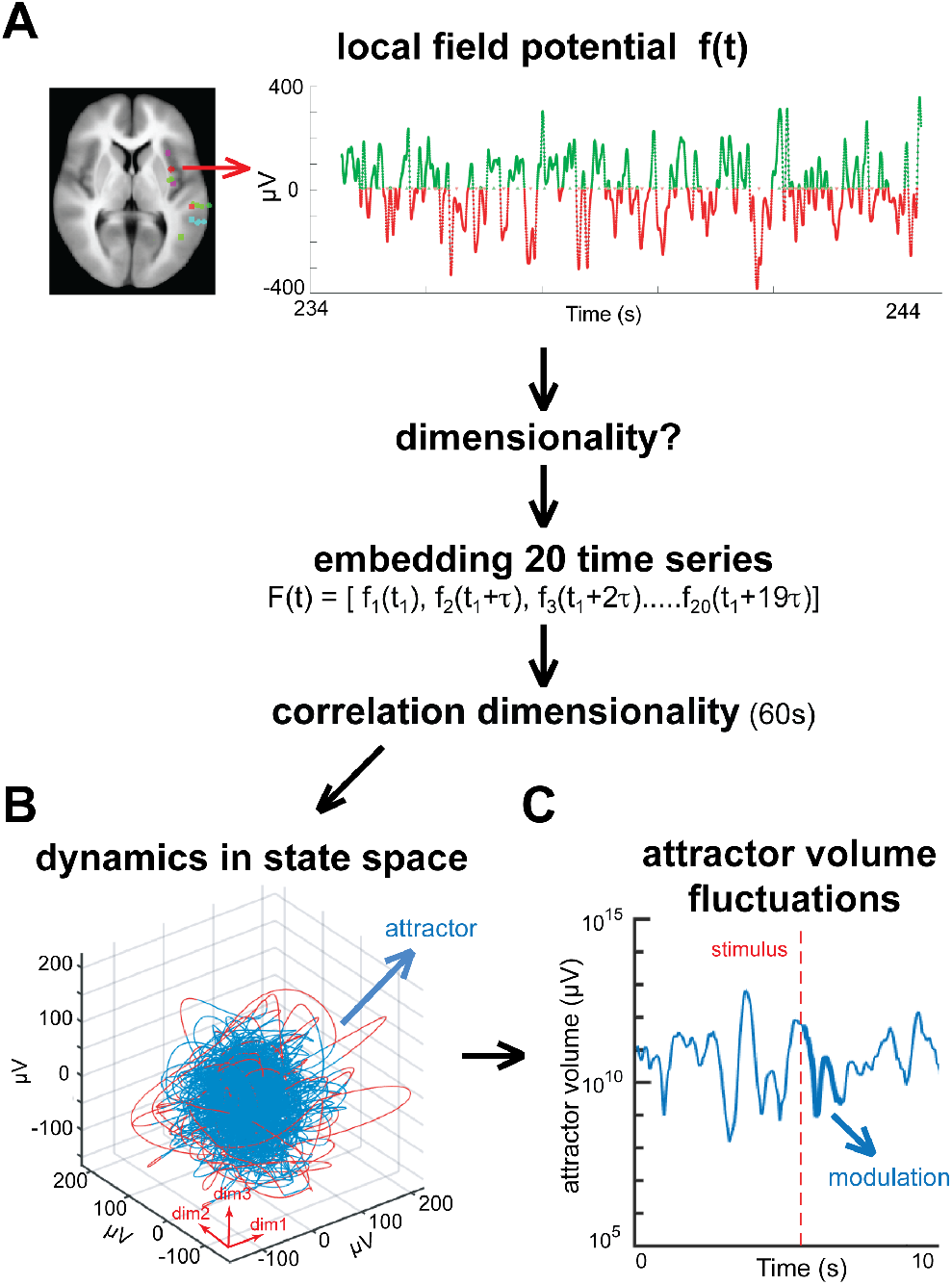
Logic of the analysis and definition of fluctuating attractor. **A**. Electrode lead with field potential *f*(*t*) sampled at 512 Hz. To find the dynamics of the local networks we must first know the *f*(*t*) dimensionality. Although the field potential is created mainly by currents from multiple dendrites of many neurons, the collective result of these many currents may be of lower dimensionality. We used embedding to estimate the dimensionality. Thereto, we constructed a vector *F*(*t*) out of 20 copies of *f*(*t*) delayed by successive multiples of τ ms. The dimensionality of *F*(*t*) is the number of dimensions needed to achieve an exhaustive description of *F*(*t*). We then calculated the correlation dimensionality from this 20-dimensional vector *F*(*t*). **B**. If, for example, the estimated dimensionality of *F*(*t*) is 3.6 we can get a good picture of the evolution of local cortical states in a 3-dimensional state space (in reality the dimensionality ranged from 3.5 to 9). In state space, each point is one local cortical state. We followed the progress of local cortical states every 1.95 ms (1/512 s) forming the blue and red parts of the trajectory. In each local network, one attractor confines the states for each behavioral condition to one hyper-elliptic part of state space. **C**. The hyper-ellipsoid expands and contracts, making the volume to enlarge and diminish. We call this a fluctuating expanding and contracting attractor (FECAT).

**Figure 2.**
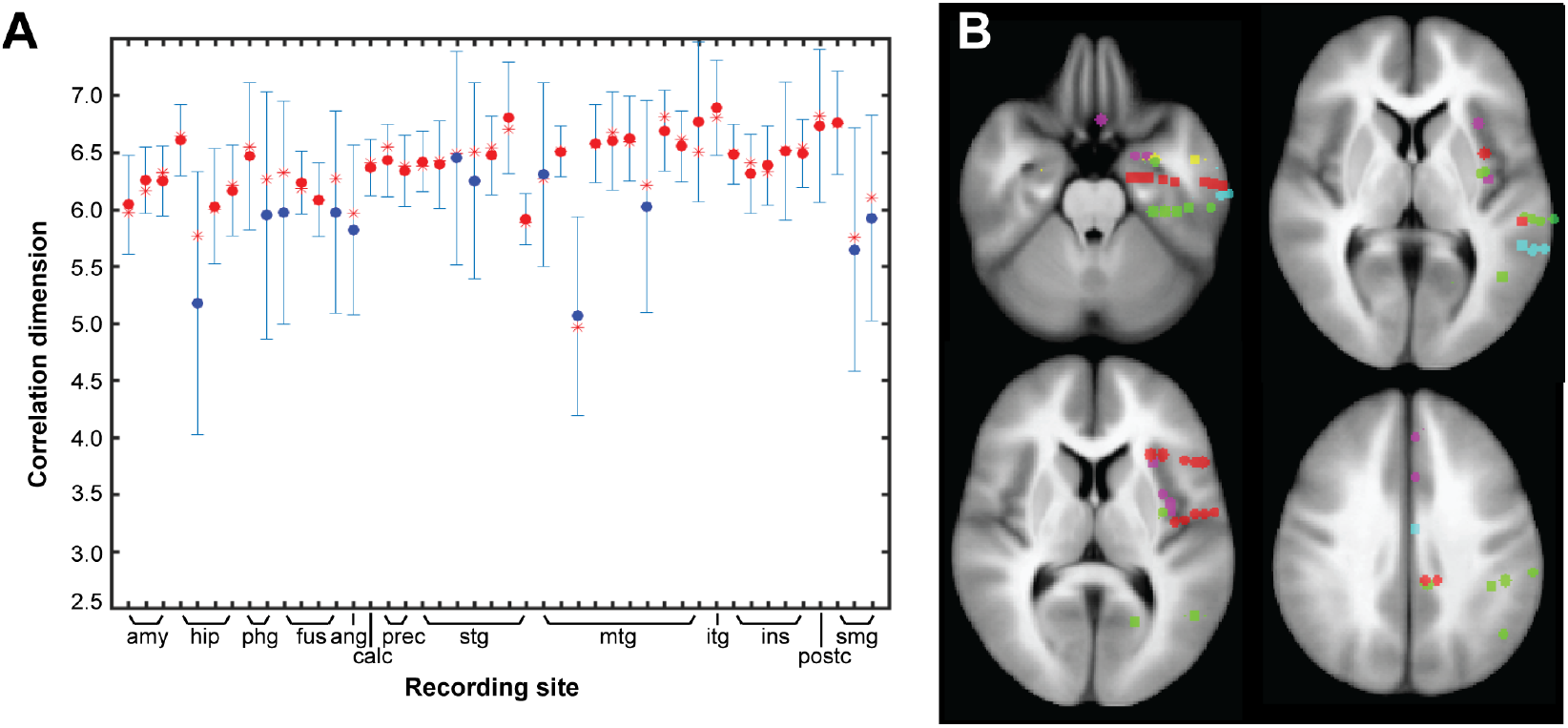
Correlation dimensionalities of field potentials. **A**. Correlation dimensions as function of anatomical location for all tests of patient 17. Red star: median. Red or blue dots: mean, depending on membership of cluster (**Figures S2 and S3)**. Standard deviation shown as error bars. Abbreviations: amy: amygdala; hip: hippocampus; phg: parahippocampal gyrus; fus: fusiform gyrus; ang: angular gyrus; calc: calcarine cortex; prec: precuneus; stg: superior temporal gyrus; mtg: middle temporal gyrus; itg: inferior temporal gyrus; ins: insula; postc: postcentral gyrus; smg: supramarginal gyrus. **B**. Axial slices of the standard brain, showing positions of electrode leads. Green spots for leads from patient 17; other colors refer to other patients.

The local correlation dimensionalities in awake conditions fell into two clusters, one with relatively high means and low standard deviations, the other with lower means and higher standard deviations (**Figures 2A and S2**). The validity of the clustering is apparent from the silhouette plots (**Methods**) in **Figure S3**.

The difference between the two clusters was also tested by traditional statistical tests (**Methods**) and was significant (p< 0.02 or less, except for patient 11; **Table S1)**. High dimensionality is associated with high complexity of dynamics and low dimensionality with lower complexity (Roschke and Basar, 1990; Stam, 1996). The estimated dimensionality for each condition and electrode lead was used to build a state space of that dimensionality, in which we studied the cortical states evolving from the local cortical network (**Figure 1; Methods**).

### Fluctuations of the volumes of cortical states

Figure 3. shows how the local cortical networks confine their evolving cortical states to a single region, shaped like a hyper-ellipsoid in state space. All cortical states in all patients (3476 60s-epochs, distributed over 290 leads, giving a total of 106782720 states) were located inside the enclosing volume of the hyper-ellipsoid in **Figure 3A**. We found qualitatively identical, albeit smaller, hyper-ellipsoids at all electrode leads and under all experimental conditions (**Methods**). Inside each hyper-ellipsoid, the temporal succession of states formed a trajectory (**Figures 3B and S4**). Thus importantly, the local dynamics measured at every cortical location and under every condition was restricted to a single hyper-elliptic region of its state space.

Data for 60 s or more are necessary to compute a reliable estimate of the correlation dimensionality (**Figure S1**), while the field potential cannot be regarded as stationary for so long. There is then a need for evaluating the dynamics at shorter, physiologically and behaviorally relevant, timescales. Consequently, we made a principal component analysis of the embedded trajectory of appropriate dimensionality. Then we estimated the volume of successive local states accumulated over 50 ms with a sliding time window of 250 ms (for choice of parameters see **Methods**). The cortical states in such short time intervals form short trajectories, rather than hyper-ellipsoids (**Figures 3 and S4**). To follow fast changes in the local states at each lead and condition, we estimated these state space changes as volumes of hyper-rectangles, **Methods)**.

**Figure 3.**
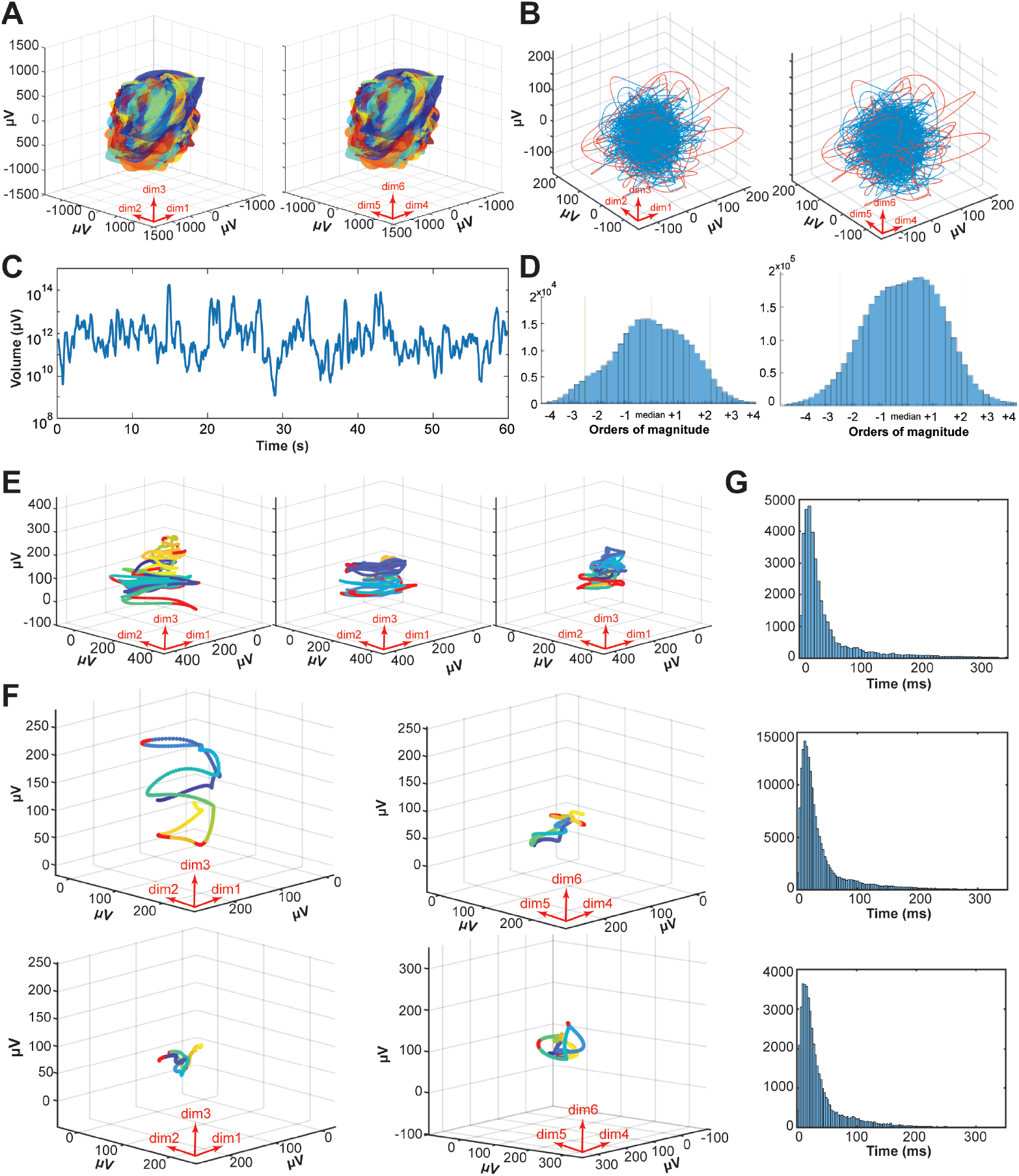
Each local cortical network constrains dynamics to one fluctuating expanding and contracting attractor. **A**. State space containing all normal states (n= 106,782,720) in all patients and conditions giving 3476 enclosing volumes (one color for each electrode-condition combination, but only the largest is visible; **STAR Methods**). Together, all enclosing volumes occupy a single, limited part of state space in shape of a hyper-ellipsoid. **B**. Single condition hyper-ellipsoid. Six embedding dimensions of **F**(t) from the posterior part of the superior temporal sulcus, reconstructed from 60 s trajectory during identification of emotional faces (patient 5, lead 24). Left: projections of the first 3 dimensions (dim). Right: dimensions 4,5,6. Red: trajectory temporarily exploiting nearby state space. Blue: trajectory inside the state space defined by the hyper-ellipsoid enclosing volume V_95%_ (**Methods**). **C**. Fluctuations of the volume calculated by equation 4 (**Methods**) of local cortical states (attractor states) during 60 s of rest. Note log_10_ scale on the y-axis. **D**. Distribution of volumes (equation 4) of fluctuating attractor. X-axis log_10_V (orders of magnitude) aligned to the median. Left: patient 12 during 20 randomly selected 60 s episodes of awake spontaneous activities. Right: all experimental conditions in awake patients (n=3363). **E**. Projections of first 3 dimensions of 5s trajectories from the parietal operculum during classifying nouns (patient 17). Fluctuating hyper-elliptic attractor of big, medium and small volume (left, center and right, respectively). Sections of trajectory exceeding Mahalanobis distances > 90% from the center of gravity are shown in red. Start and end of 5 s episode trajectory are shown in dark blue and yellow, respectively. **F**. Projections of first 3 dimensions of 1 s trajectories from fusiform gyrus during recall of visual events at admission day (patient 12). Left: big volume trajectory of dimensions 1-3 (upper part) and 4-6 (lower part); Right: small volume trajectory. Conventions as in E. **G**. Distributions (log normal, **Figure S4**) of times to return for outside states to the enclosing V_90%_, when V_90%_ is big (top), medium (middle) and small (bottom).

Note that we use two complementary methods to estimate two different volumes of the local cortical states. The hyper-ellipsoid method was used for successions of cortical states 1s and longer (**Figure 3A, B, E, F**). The hyper-rectangle method was used for shorter successions of cortical states (equation 4, **Methods**; **Figures 3C and 3D**). During ongoing autonomous activity only (rest condition), the enclosing hyper-rectangle volume continuously expanded and contracted (**Figure 3C**). In all (n=110) measurements of the rest condition, the enclosing volume continuously expanded and contracted irregularly. These expansions and contractions were also the dominant result at all cortical sites and under all tests and conditions (n=3476) (**Figures 3D and S5; database)**. For comparison, we also plotted the similar volume fluctuations from 20 randomly selected 60 s episodes during spontaneous behavior of one awake patient, i.e. in addition to the experimental conditions. In all cases, the ranges of the enclosing volume (from minimum to maximum) were at least four orders of magnitude, with no difference in the ranges between spontaneous behavior and experimental conditions (**Figure 3D** right and left). Fluctuations of the (one-dimensional) cortical field potential are well known (Einevoll et al., 2014; Pesaran et al., 2018). However, that these vigorous fluctuations should be the dominant feature of the dynamics of evolving cortical states at full dimensionality was not foreseen.

Over shorter (1s, 5s) time scales, the flow of states can be localized to well-defined trajectories, as shown in the projections in **Figures 3E and 3F**. The volume fluctuations were irregular (**Figure S5; database**) meaning that in each cortical network, a shorter succession of cortical states expanded and contracted irregularly. Equivalently, one can describe the expansion and contraction as stretching and contraction of the multidimensional trajectory forming the succession of states. These expansions and contractions continued under all conditions, with no exceptions (3476 out of n= 3476) **(database**). These volume fluctuations were not very fast (50 ms to a few hundreds of ms)(for details and flow of states, see **Methods**).

### Attractor properties of the dynamics

Fast volume fluctuations of four orders of magnitude indicate that the local networks they originate from produce unstable dynamics. For this reason, we examined the stability of the evolving cortical states. In **Figure 3B** we set the enclosing volume as the volume of the hyper-ellipsoid enclosing all states whose Mahalanobis distances from the center lie inside the 95% quantile (**Methods**). As the outside states quickly returned to the 95% enclosing volume, we suspected that this observed dynamic could be attractor-like. Therefore, we examined the fate of outside states at all time scales up to 60 s. In addition, we examined outside states when the enclosing volume of the putative attractor, because of its fluctuations, was small, medium and large (for definitions see **Methods**). We defined the putative attractor as the set of states whose Mahalanobis distances from the center of the enclosing hyper-elliptic volume were less than 90% of the maximum measured value. Thus, large volumes, medium-sized volumes and small volumes all had by definition 10% of states as outliers (**Figures 3E and 3F**). Most of these states returned to the putative attractor within 50 ms, at a different position than that from which they had departed, as is typical for a dynamical system (**Figures 3G and S4**). Changing the definition of the outside states to 20% or 5% did not change the distribution of the return times (**Fig. S4**, see also **Fig. S4**, for other proofs of attractor properties). The states evolved within a hyper-elliptic region in state space. States which occasionally escaped this region were swiftly pulled back by the attractor, irrespective of the size of the hyper-ellipse.

To sum up, each local cortical network produces local cortical states. The succession of states forms a trajectory which for 90% is contained inside a hyper-elliptic object. The states inside the object attracts states outside and pulls the outside states back. Therefore, the object has attractor properties. This attractor expands and contracts continuously in state space, thereby expanding and compressing the states it contains. Or, phrased equivalently, stretches and contracts the trajectory it contains (**Figures 3E and 3F**).

These results are compatible with single attractor dynamics. This analysis, however, does not distinguish whether what we defined as outside states in reality were parts of the attractor itself - or departures from the attractor that were rapidly pulled back. None of our experimental conditions succeeded to perturb the attractor systematically. This shows the overall stability in of the local cortical networks under normal conditions. Moreover, the state flows were fast in all dimensions within the state space enclosed by the attractor. This minimizes the likelihood for alternative dynamics hidden in the field potentials (**Figure S6**). Furthermore, an attractor, as normally defined, has a single invariant dimensionality and occupies an invariant part of state space. What we observed is not an attractor in the normal sense, because of the expansions and contractions in state space. To describe this lower-dimensional fluctuating expanding and contracting attracting object, we coined the term fluctuating expanding and contracting attractor (FECAT).

### Fluctuating expanding contracting attractor dynamic is a universal large-scale cortical state dynamic

In these 3476 episodes, comprising all tested conditions and all 290 leads we could not find any exception from fluctuating attractor dynamic (**Figures 3A, 3B, 3G, S4** and **S6; database**).

This means that local networks from different cortical locations and within different cortical areas used the same type of dynamic to drive their cortical states. Indeed, local cortical networks in prefrontal cortex, primary visual cortex, supramarginal gyrus, and hippocampus, have widely different anatomical and neurotransmitter receptor architectures (Elston, 2003; Zilles and Amunts, 2010). Notwithstanding these architectonic differences, the local networks created cortical states using FECAT-dynamic (**Figure 4; database)**.

**Figure 4.**
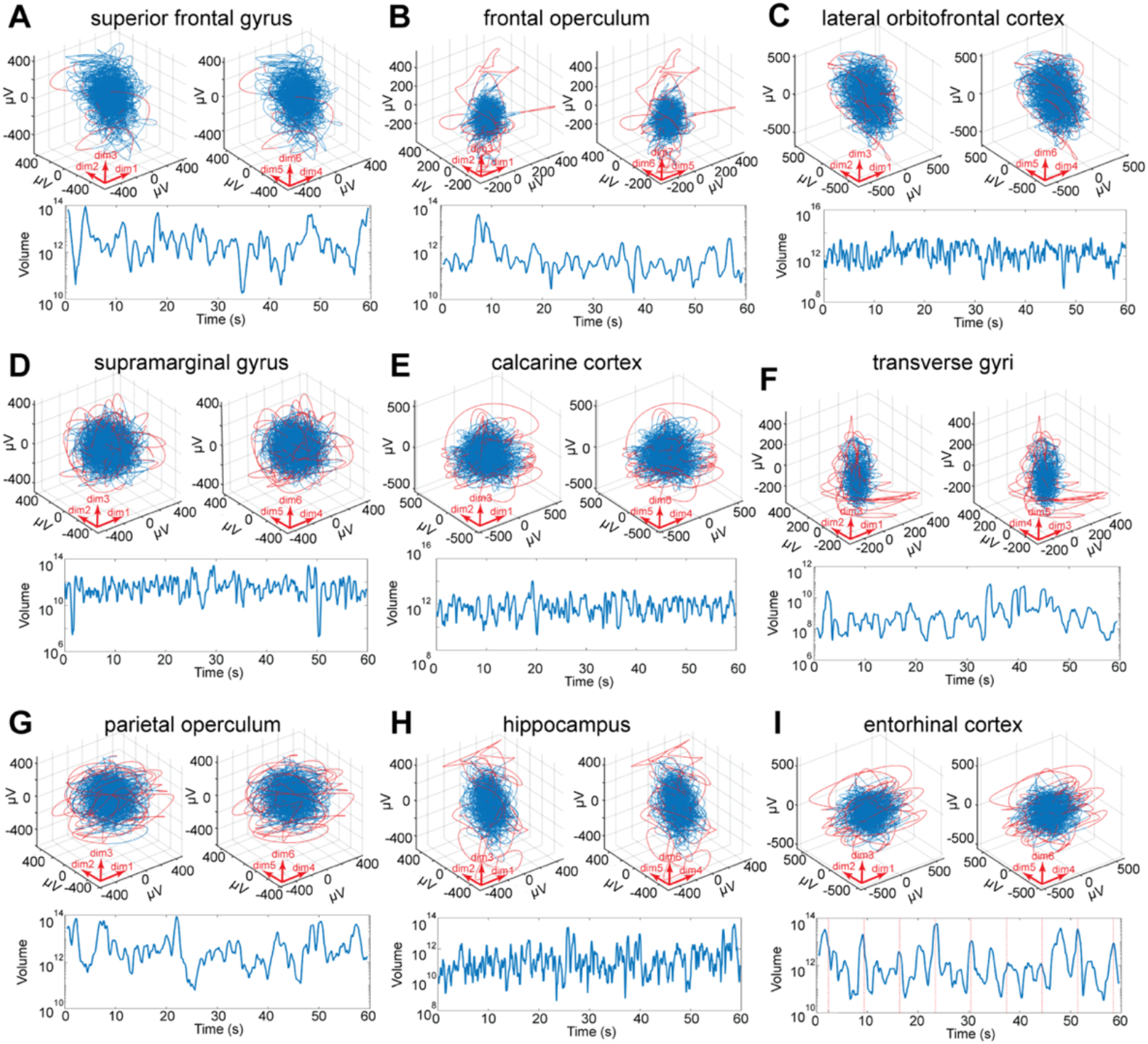
Different cortical network architectures create cortical states with FECAT-dynamic. Projections of attractor trajectories over 60 s; Red curves: trajectories temporarily exploiting nearby state space. Blue curves: trajectories inside the state space defined by the enclosing hyper-elliptic volume V_95%_. Below: fluctuations of the volume of cortical states (equation 4, **STAR Methods**) during the same 60 s. **A**. Stereognosis (patient 6). **B**. Conversation (patient 4). **C**. Ortho-nasal smell (patient 16). **D**. Motor test 6 (patient 17). **E**. Visual recognition (patient 17). **F**. Memory retrieval, mental navigation (patient 5). **G**. Watching TV (patient 12). **H**. Eating (patient 14). **I**. Facial emotion identification (patient 4).

Similarly, during all tested conditions, cortical states evolved under the rule of FECAT-dynamic. For example, cortical states during spontaneous behavior, rest condition and during different brain functions such as mental navigation, classification of nouns, somatosensory attention, tasting sweets etc., were all products of FECAT-dynamic (**Figure 5; database**). The different conditions and brain functions produced somewhat different patterns of expansions and contractions (**Figures 5 and S5, database)**. However, single trial tests were associated with a single expansion and contraction.

**Figure 5.**
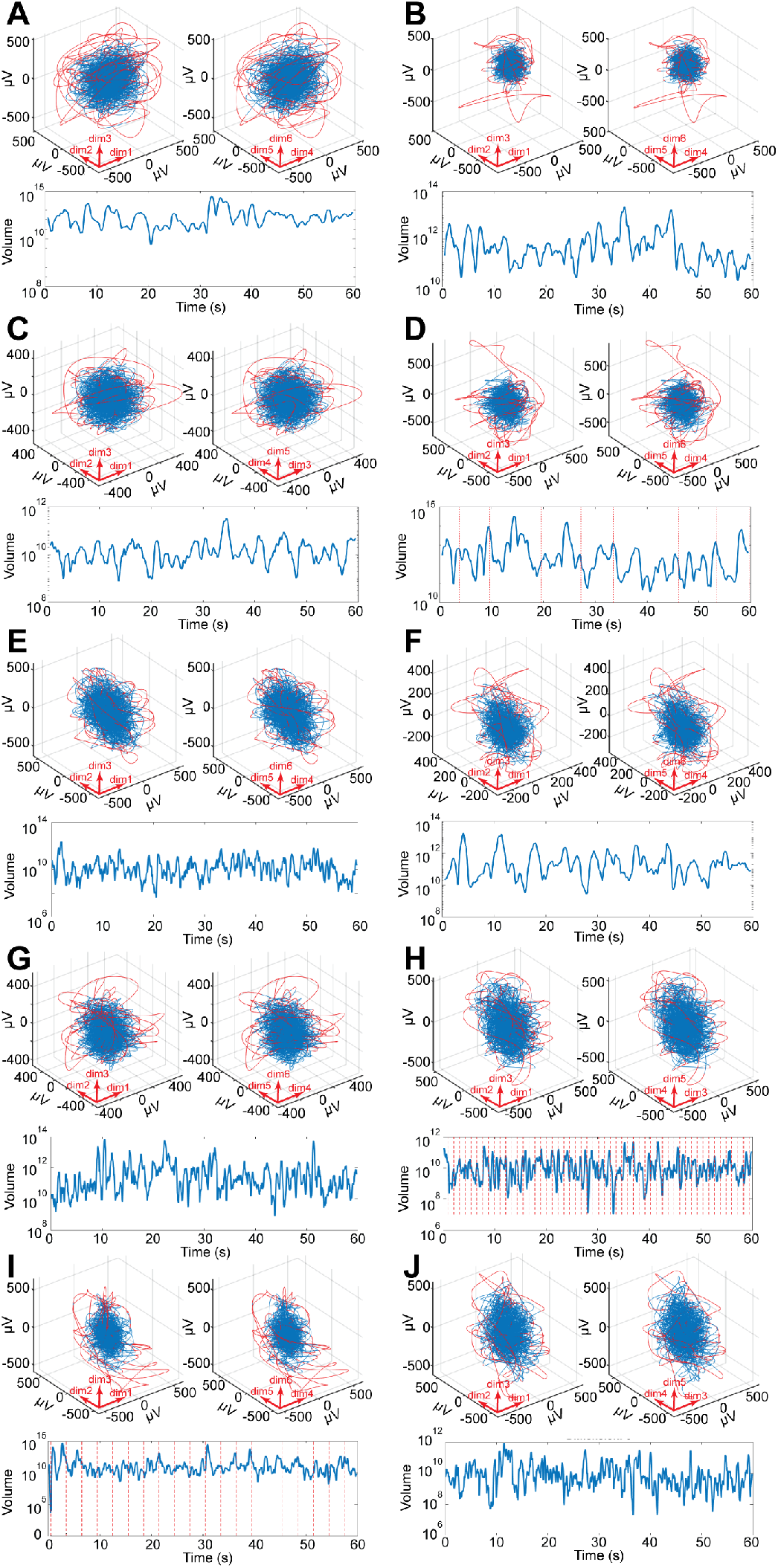
Different brain functions emerging from FECAT dynamic. Projections of attractor trajectories over 60s; Red curves: trajectories temporarily exploiting nearby state space. Blue curves: trajectories inside the state space defined by the enclosing hyper-elliptic volume V_95%_ (**STAR Methods**). Below: fluctuations of the volume of cortical states (equation 4, **STAR Methods**) during the same 60 s. **A**. Control (rest) condition, superior temporal gyrus, anterior part (patient 4). **B**. Mental navigation (memory retrieval test 8), posterior hippocampus (patient 6). **C**. Memory retrieval test 10 (admission day), frontal operculum (patient 12). **D**. Classification of nouns, parahippocampal gyrus (patient 8). **E**. Somatosensory attention, anterior insula (patient 16). **F**. Watching TV, superior frontal gyrus, anterior part (patient 9). **G**. Visual motion to right (visual test 7), posterior hippocampus (patient 15). **H**. Visual test 4, middle part of inferior frontal gyrus (patient 14). Note entrainment. **I**. Recognition (visual test 6), subiculum (patient 10). **J**. Taste, amygdala (patient 16)

### Attractor size fluctuations reshape to single trial input

Single trial tasks reshaped the ongoing expansions and contractions, but did not perturb the dynamic. The volume fluctuations of the attractor thus changed to single trial input, for example, asking to imagine the face of a well-known person. This elicited a small expansion followed by a slightly longer contraction time-locked to the invitation to imagine and thus modifying the ongoing fluctuations (**Figure 6A and 6B)**. Similarly, other single trial tasks in which patients were asked to identify facial emotional expressions, imagine well-known landmarks or tourist attractions, judge whether spoken nouns were concrete or abstract, look at geometrical shapes, and recognize natural scenes also elicited a small expansion followed by a contraction concurring with the imagery, problem solving and perception (**Figures 5, 6F and S5**). Each patient did from 2 to 7 such single trial tests. This shows that a single trial engagement was associated with an expansion followed by a contraction of the volume of the fluctuating attractor. This is compatible also with the stretching and compression of the trajectories in **Figures 3E and 3F**.

**Figure 6.**
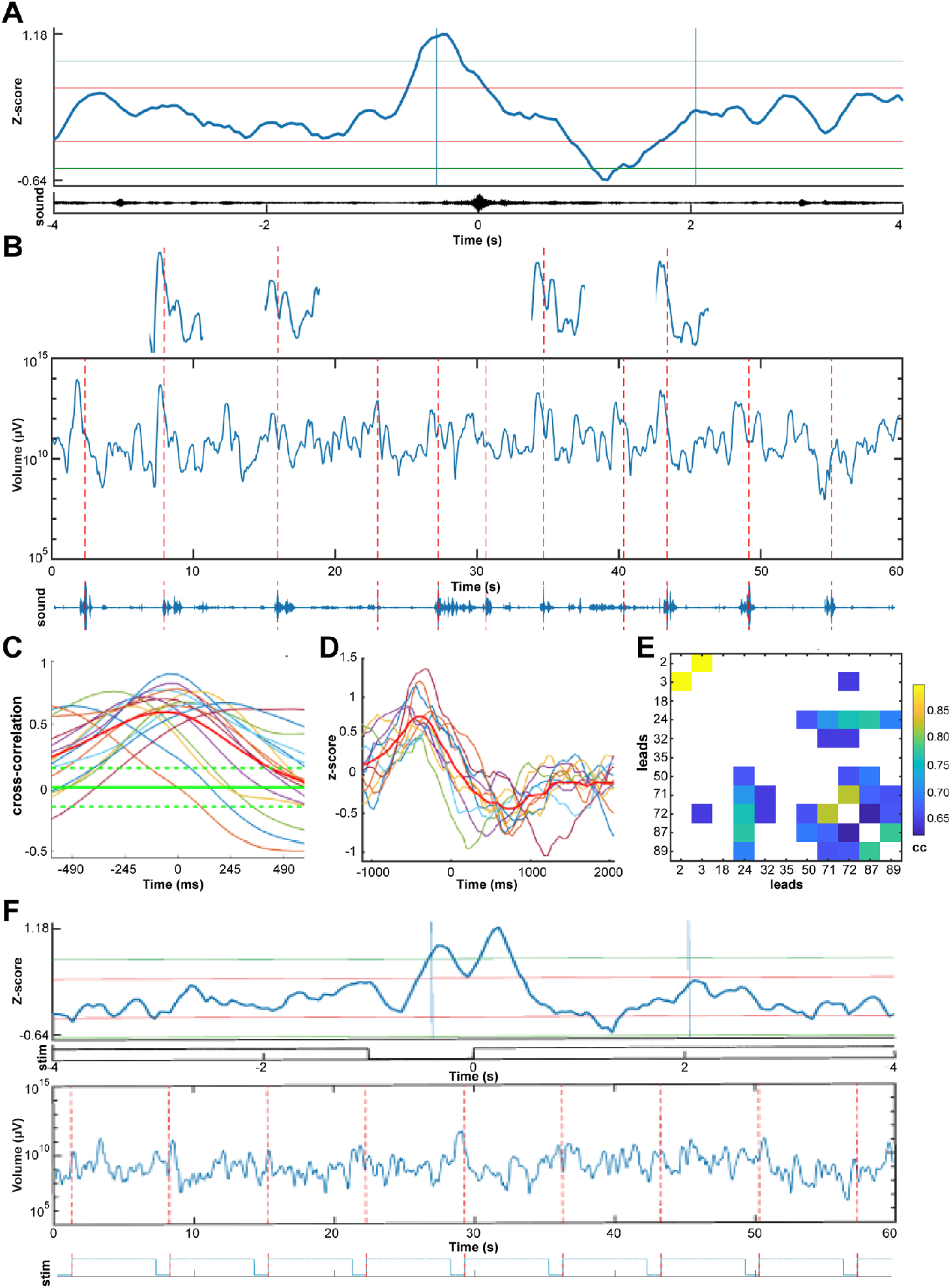
Single trial expansion and contraction of the fluctuating attractor. **A**. 4 s average of 11 z-scores of attractor volume (V) showing one expansion and contraction. Average of the sound track below. Each trial aligned to the peak of the sound. **B**. Fluctuations in 11 trials of V of the attractor in log_10_ scale, recall of faces (memory retrieval test 1, **Methods**). Patient 17, lead 50, cortex in posterior part of the superior temporal sulcus; below: sound track with peaks indicated by the red stippled lines for each of the names of the 11 well-known persons to imagine (**Methods**). **C**. Statistically significant cross-correlations of average z-scores between leads engaged in the task. Green stippled lines: 95% confidence limits. **D**. z-score averages, with mean in red, of expansions and contractions from the leads engaged in the task. **E**. Pairwise cross-correlations (cc) of the leads in D. **F**. Top: z-score average of V from 9 trials; below: average photodiode signal showing end of last screen presentation, 1 s black screen interruption, and first 4 s of emotional expression to judge. Bottom: fluctuating volume of the attractor in log_10_ scale with photodiode signal of 9 single trial screen presentations below. Patient. 5, lead 6, 6 dimensions, middle temporal gyrus.

But how do the cortical networks work together to perform these brain functions? The face imagery test (**Figures 6A and 6B**) was associated with such an induced expansion and contraction in many other local networks. In these local networks, these modified expansions and contractions of the fluctuating attractors cross-correlated mutually and cross-correlated with the modified expansion and contraction of the local cortical states in **Figure 6A**. These pairwise cross-correlations appeared with different lags (**Figures 6C and 6D**). However, not all local cortical networks had significant cross-correlations of the fluctuations (**Figure S5**). Thus, only a subset of the many local networks modified and correlated their attractor expansions and contractions during the test, while the remaining networks continued their un-correlated fluctuations (**Figure S5**).

These results also held for the other single-trial tests. The local networks engaged by cross-correlating their single expansion-contraction with different lags. With few exceptions, each of these tests was associated with a unique combination of cross-correlated local cortical networks (**Table S2**). On average, 43.5 ± 24.8 % (SD) of the local networks engaged with cross-correlated fluctuations in these tests (n=43). This demonstrates the sensitivity of the attractor volume fluctuations to single transients and stimuli. At the macroscopic scale, the single trial tests seem to be operated by different combinations of cortical local networks.

We also observed expansions and contractions comparable to those in **Figure 6** in the absence of sensory stimulation (example in **Figure 3F)**. All expansions and contractions of attractor volumes during all conditions can be found in the **database** (https://github.com/ChristofferKKHansen/RolandLab).

### Entrainment of cortical states

Rhythmic visual stimulation entrains the volume fluctuations of the attractors. Entrainment of field potentials in visual cortex of monkeys has been reported (Lakatos et al. 2008). In visual tests 4,5 and 11, patients were instructed to watch the items shown on the monitor, without responding. This elicited an entrainment of the expansions and contractions of the attractor, with a frequency identical to (**Figures 7A and 7F**) or (occasionally) double that of the stimulus presentation rate (**Figure 7G**). Rhythmic visual presentations with no intermission of a black screen (visual test 3), and stimulus durations of 6 s (facial emotions) did not produce entrainment. In contrast, regular presentation of visual stimuli of 125 or 1000 ms duration at 0.5 or 1Hz did produce entrainment (including visual test 6).

**Figure 7.**
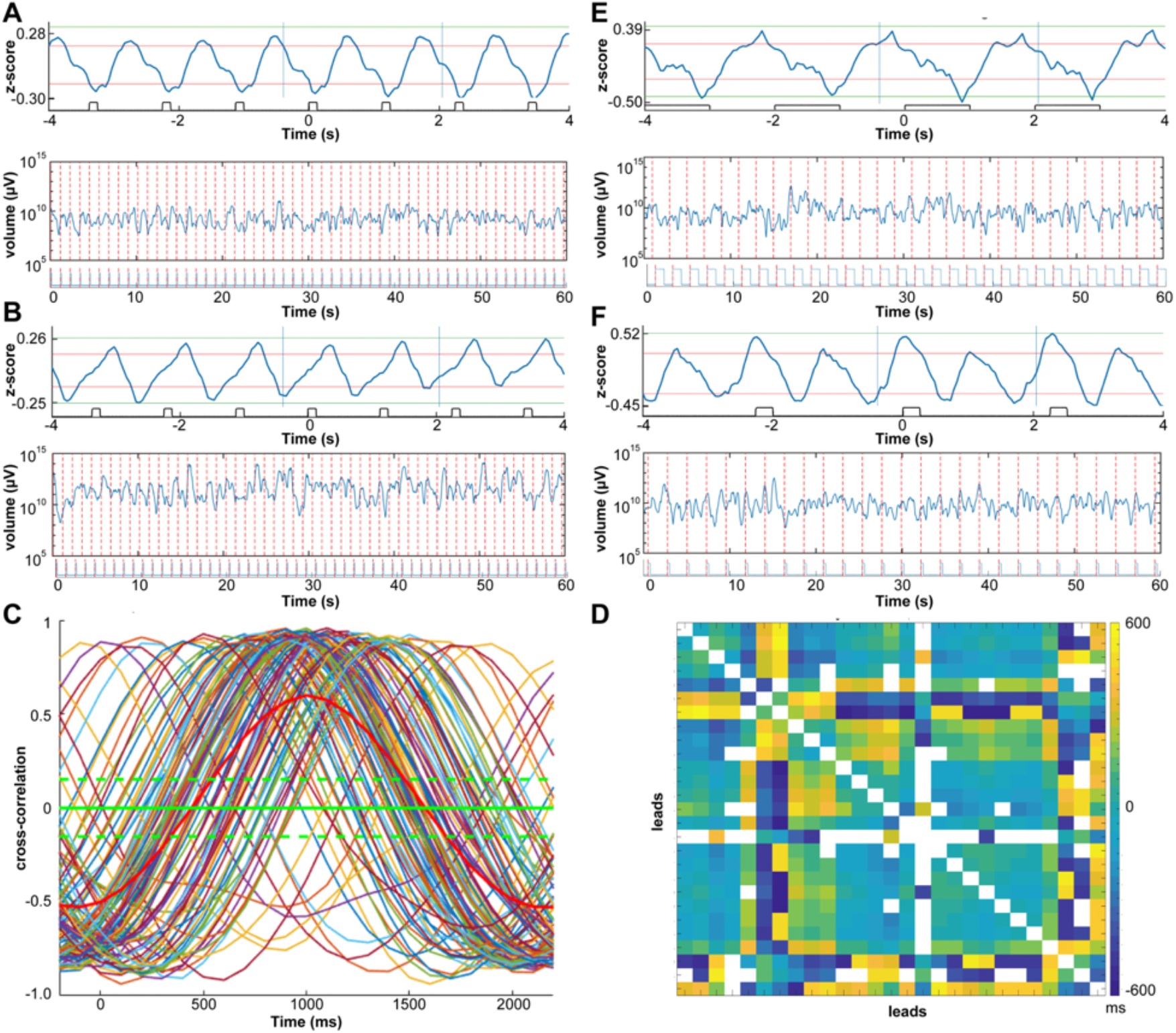
Entrainment of fluctuating attractors. **A**. Average z-scores of volume fluctuations over trials with average stimulus presentation. Bottom: 60 s episode of fluctuating size V of the attractor in log_10_ scale with onset of trials marked with red stippled lines. Natural pictures presented 125 ms with 1 s black screen intermissions (visual test 4). Patient 11 lead 2, anterior part of cingulate gyrus. **B**. Same patient, same test, but lead 101, middle temporal gyrus, posterior part; phase shift of the fluctuating expansions-contractions of the attractor (compared to A). **C**. Pairwise cross-correlations above 70% quantile of average z-scores during the task for the time segment defined by the 2 vertical lines in A. Green stippled lines: 95% confidence limits. **D**. Matrix of the pairwise lags (in ms) for all leads. **E**. Visual recognition test, patient 6, lead 2, anterior part of cingulate cortex. Average of z-scores (top), 60 s episode of recognition of fluctuating size V of the attractor in log10 scale with photodiode signal of picture presentation below. **F**. Fluctuating attractor entrainment of double frequency. Geometrical pictures (visual test 11) patient 5, lead 25, cortex lining posterior part of superior temporal sulcus.

As for the other single trial tests, one stimulus elicited one expansion followed by one contraction (**Figures 7A, 7B, 7E and 7F**). The phase of the expansion-contraction oscillation however varied with cortical location. In some local networks, the expansion led the stimulus presentation (example in **Figures 7A and 7E**), while at other sites it trailed it (**Figures 7B and 7F**). The frequency of these entrained oscillations was strictly locked to the presentation frequency (n= 41 out of 41 tests) (**Figures 7A, 7B, 7E and 7F**) with lags ranging from -500 to + 500 ms (**Figures 7C and 7D)**. In these strictly rhythmic trials, anticipation of the next stimulus as well as evoked activity may explain the large range in lags (see also **Figure 6F**) (Gillies et al., 2019; Staresina et al., 2019).

The proportion of leads showing attractor volume entrainment in the four visual tests (4, 5, 6 and 11; **Methods**) ranged from 0% to 100% (**Table S2**). Natural pictures presented for 125 ms at 1 Hz engaged from 91 % to 100% of the leads (mean 97%; n=8). The entrainment was not obvious in the raw data (**Fig. 5H**), but it was evident in the stimulus onset time-locked averaged data (**Figures 7A, 7B, 7E and 7F**). This shows that the system is not simply undergoing oscillatory (limit-cycle) dynamics; the time-locked oscillations here are just another property of the attractor fluctuations.

## DISCUSSION

Our results provide an experimentally-based overall explanation of how cortical states contained in field potentials from the human cerebral cortex evolve during a wide spectrum of behavioral conditions. All examined local cortical networks produced successions of local cortical states forming trajectories that stretched and contracted irregularly and unceasingly. The trajectories predominantly resided inside a single multidimensional object, an attractor that continuously expanded and contracted in state space. The states inside this attractor pulled states that temporarily escaped back into the attractor, irrespective of the volume of the attractor. Stimuli, single trial tasks, and other behavioral conditions temporarily reshaped the expansions and contractions and altered the progress velocity of cortical states, but did not radically alter or change this dynamic into other dynamical regimen. The reshaped expansions and contractions cross-correlated between selected subsets of cortical networks, depending on the test. This hitherto unreported fluctuating expanding and contracting attractor dynamic gives a simple general explanation of larger scale (postsynaptic) dynamics of the cerebral cortex at behaviorally relevant time-scales.

By extracting non-linear field potential dynamics in state space of sufficiently high dimensionality (Takens 1981) and by avoiding methods requiring stationarity, we discovered the fluctuating attractors. A low-dimensional, slowly decaying, oscillating spiral attractor was recently described (Bruno et al. 2017). However, we were unable to find attractors similar to FECAT in the literature. Attractors of similar dimensionalities to FECAT derived from the human EEG have been proposed (Babloyantz and Destexhe, 1986; Roschke and Basar, 1990, Stam, 1996) based on a working assumption of stationarity. It is possible that these putative EEG attractors are in fact FECAT attractors, which could be tested, for example by the strategy outlined in **Fig. 1** and **Methods**.

The state space dimensionalities in our data at all recorded sites were larger than 3 but less than 10. Previous studies on cortical dynamics reported dimensionalities up to 3 (Tognioli and Scott Kelso, 2011; Knierim and Zhang, 2012; Rabinovich et al., 2012; Priesemann et al., 2013; Wimmer et al., 2014; Oprisan et al., 2015; Bruno et al., 2017; Hennequin et al., 2018). One reason for this discrepancy might be that FECAT-dynamic was derived from the actual cortical field potential signal, instead of being a theoretical attractor model borrowed from physics or other disciplines.

The electrode lead sites were not uniformly distributed over the cortex in the whole material, and were sparsely scattered in individual patients. This precluded a coherent anatomical reconstruction of the lead positions for examining systematic relations between tasks, anatomical localization and the expansions and contractions and their spatio-temporal cortical dynamics (Arieli et al., 1995; Ferezou et al., 2007; Ayzenshtat et al., 2010; Roland et al., 2017; Muller et al., 2018).

### Evolution of postsynaptic cortical states and local excitation and inhibition

Local networks in prefrontal, parietal and temporal cortices, hippocampus and amygdala all operated with FECAT-dynamic. These networks have well-described different anatomical and receptor architecture (Zilles and Amunts, 2010). The common FECAT-dynamic thus should have a common cause. One explanation of this consistent result is that delays and timing of excitation and inhibition are preserved in the field potentials from these different networks and expressed in the FECAT-dynamic.

The field potential is a weighted sum of the extracellular currents in the neighborhood of the electrode lead. The main sources of these extracellular currents are the underlying multiple postsynaptic membrane currents (Einevoll et al., 2014; Pesaran et al., 2018). The field potential is measured at a relatively coarse mm^3^ scale, causing many details of the biophysical computations of inward and outward currents to get lost (Icaruso et al., 2017; McCormick et al., 2020). Only closely timed and dominant excitations and inhibitions in the network may remain in the multidimensional field potential. So, it might well be that architectural differences in the local networks are responsible for important biophysical computations that are not captured at the mm^3^ scale. At this scale, the expansions, contractions, progress of states and attraction could be caused by multidimensional action of excitation and inhibition in the local network. Without excitation, states will not progress and expand in state space. Without inhibition, evolving states will not contract, and states escaping from the attractor will not be pulled back (Roland et al., 2017).

The field potential is closely correlated and in phase with the voltage sensitive dye signal in the cerebral cortex of rodents, carnivores, and awake primates (Arieli et al. 1995; Ferezou et al. 2007; Lippert et al. 2007; Xu et al. 2007; Ayzenshtat et al. 2010). The visual cortical areas in the ferret have similar short stretching of trajectories in state space when excitation dominates the local network, followed by a slower 1 s contraction when inhibition dominates (Roland et al. 2017). Thus, the stimulus-induced expansions and contractions of cortical states we find in human local networks may relate to local dominance and timing of excitation, followed by local dominance and timing of inhibition. The ongoing expansions and contractions in state space observed in the absence of experimental stimuli may also reflect such shifts.

### Implications of FECAT-dynamic

An advantage of such an organization of common network dynamics is that it facilitates the communication between different local networks. If local networks operated with different dynamics, it would take time to mutually adjust the dynamics. Indeed, in the single trial tests, the reshaped expansion and contraction of cortical states cross-correlated between a subset of cortical networks. What is communicated between the networks must be reflected in the timing and dominance of excitation and inhibition in the target networks. These excitation-inhibition shifts exist at multiple (local spatial) scales. Finer details in perception, imagery and motor control which cannot be resolved at this mm^3^ scale could be resolved by mechanisms also affecting the delayed timing of excitation and inhibition at microscopic scales (Harvey and Roland, 2013; Dehghani et al., 2016; McCormick et al., 2020). The consistent FECAT-dynamic is the overriding principle at the mm^3^ scale. However, whether FECAT-dynamic is preserved at finer scales and how the state expansions and contractions relate to local excitation and inhibition are questions that can be tested more readily in animals.

If the multi-dimensional timing and dominance between excitation and inhibition is expressed in the FECAT-dynamic, this would also explain how all the different brain functions we examined could emerge from FECAT-dynamic. Having only one (type of) fluctuating attractor dynamic operating under all conditions means that the cortex does not need to change dynamics to react to new transients and to shift to another brain function.

Changes in field potentials that are not locked to stimulus presentation or motor behavior are often regarded as background activity, obscure ongoing activity, default mode, resting, “dark activity” or noise (Buzsaki, 2019; Singer et al., 2019; but see McCormick et al., 2020). However, cortical states in all the different behavioral conditions, spontaneous and resting states included, evolved under FECAT-dynamic. Also, the expansions and contractions of cortical states during experimental conditions were within the same ranges as during spontaneous activities and rest (**Figures 3C, 3D and 3G**). These results show that there is no reason to maintain that spontaneous attractor state fluctuations during rest and under spontaneous activity in the awake conditions are noise or driven by other types of dynamics than the dynamic driving perception, retrieval of memories, thinking, and preparation of motor activity. The FECAT-dynamic accounting for all these conditions ensures that cortical networks easily switch from internally initiated functions to externally initiated functions and vice versa. Thus, spontaneous ongoing and resting activity, sensory evoked and preparatory motor activity is unified by the FECAT-dynamic creating local cortical states.

It is therefore unlikely that FECAT-dynamic is caused by stimuli, test demands and other external causes. The progress, the expansions and contractions, and attractor dynamics continued without external signals or demands. This result, and the finding that all conditions evolved under FECAT-dynamic, indicate that FECAT is an ongoing continuous intrinsic dynamic of cortical local networks that can be modified, but not changed or replaced by another dynamic in response to external demands or stimuli. Having local networks with ever ongoing progress, expansions and contractions of cortical attractor states that can be modified by other cortical networks has important biological advantages. It facilitates the communication between local networks and eliminates needs for starting mechanisms or stimuli initiating cortical operations.

Cortical networks must remain permanently stable, preventing runaway excitation (Rabinovich et al., 2012; Breakspear, 2017; Singer et al., 2019). On the other hand, most known stable attractors prevent the network to react to small transients (Tognioli and Kelso, 2011, Rabinovich et al., 2012; Wimmer et al., 2014). The attractor property of FECAT-dynamic explains how cortical states under all (physiological) conditions can evolve with stability (i.e. do not split (bifurcate), change into another dynamic or run off). Yet, the changes of the fluctuations demonstrate the sensitivity of this dynamic to small single trial inputs, as expected from a complex dynamical system. Unifying vigorous state fluctuations shaping cortical functions with attractor stability is a unique biological solution. This solution unifies cortical stability with sensitivity to small transient inputs, and resolves a long-lasting conundrum in theoretical and computational neuroscience (Tognioli and Kelso, 2011; Rabinovich et al., 2012; Breakspear, 2017; Hennequin et al., 2018).

One principal result is that irregular fluctuations of the attractor volumes are the rule. However, FECAT-dynamic demonstrates that both fluctuating states and entrained oscillatory cortical states can arise from the same dynamic. This resolves a longstanding controversy between scientists who regard asynchronous irregular cortical states as characteristic for the cerebral cortex, and others who emphasize oscillations as fundamental for cortical operations (Rheinberger and Jasper, 1937, Einevoll et al., 2014; Muller et al., 2018; Breakspear, 2017; Pesaran et al., 2018; Buzsaki, 2019; Singer et al., 2019; Ju and Basset, 2020; McCormick et al., 2020).

Like almost all other dynamics, FECAT-dynamic accounts for the *temporal* evolution of states in local cortical networks. Temporal dynamics may not suffice to explain all cognitive functions, behaviors and perceptions. The diversity of cortical functions is likely to depend also on the spatial operations within and among local cortical networks. One local cortical network is often incapable of distinguishing clear differences in stimuli, especially in single trials (Forsberg et al. 2016). Different conditions might differ by the detailed nature of the attractor fluctuations and by their *spatio-*temporal cross-correlations in cortical space. Cross-correlations of attractor volume fluctuations captured the cooperation between local cortical networks across the cortex in the examples provided. This makes single trial cortical operations robust. Notably, the expansions and contractions were cross-correlated, not the field potentials themselves as in previous reports (Muller et al., 2018; Singer et al., 2019). The results also indicated that cross-correlated expansions and contractions of cortical states among a certain combination of local networks characterized a particular single trial test. This suggests that engagements in different tasks may be distinguished by *spatio*-temporal differences in the participations of local networks (Arieli et al. 1995; Ferezou et al., 2007; Ayzenshtat et al., 2010; Iacaruso et al., 2017; Roland, 2017; Singer et al., 2019).

The trajectories inside the attractor show how cortical states evolve in state space and time. Since the trajectories of the multidimensional field potential in state space continuously stretched and contracted, our results are incompatible with dynamics temporarily settling into particular stable states (equilibria) or representations. This applied even to conditions in which patients were exposed to stationary pictures or imagined stationary objects. These conditions were associated with a short expansion, stretching the trajectory, followed by a 1 to 2 s contraction of the trajectory (**Figures 4, 5 and 6**). When the trajectory stretches, the state-space-time sequence (making the trajectory) gets stretched (evolve faster). When the trajectory contracts, the state-space-time shortens (evolve more slowly). The contractions, as argued above, may relate to an overall inhibition in the local networks coinciding with visual perception (Lee, 2003; Ozeki et al. 2009; Harvey and Roland, 2013; Roland et al., 2017). Thus, cross-correlated contractions in state-space-time between networks may relate to the subjective experience of scenes or objects, since the experience concurred with the contractions.

Cortical states derived from field potentials are traditionally associated with more global conditions such as sleep and being awake, attentive and alert, in time scales ranging from seconds to hours (Rheinberger and Jasper, 1937; Stam, 1996; Busaki, 2019; Singer et al., 2019). However, the postsynaptic currents in a local cortical network also reflect the collective computations contributing to brain functions at the mm^3^ spatial scale and the ms time scale (Arieli et al., 1995; Ferezou et al., 2007; Lippert et al., 2007; Xu et al., 2007; Ayzenshtat et al., 2010; Harvey and Roland, 2013; Roland et al. 2017). The cortical states extracted from the field potentials and their flow, stretching and contractions should therefore also reflect contributions to specific brain functions being performed.

The fluctuating expanding and contracting attractor is a mathematical object. This makes it a theoretically overall and simple description of how cortical states evolve that is easy to test experimentally, also at finer scales of observation and in other species. However, FECAT dynamic does not deal with spiking dynamics.

### Perspectives

Consistent fluctuating expanding and contracting attractor dynamic was used as an overall collective mechanism to drive evolving cortical states at the mm^3^ spatial scale. If there were no common mechanisms controlling cortical operations, the alternative is to record membrane currents from all cortical neurons simultaneously under a large variety of conditions to explain how the cortex works (BRAIN 2025). Our results explain the stability and the sensitivity of the local networks, the continuous changes of cortical states and how local cortical states evolve in operations under many natural conditions. They demonstrate that internal, evoked, and motor states under these conditions arise from the same dynamic, which could facilitate cortical operations. The mathematical structure of this dynamic makes it an explanation of cortical dynamics that can be tested experimentally. Our data left no evidence of any stationarity, stable patterns, stable states or equilibria or other alternative types of dynamics. Our results are in accordance with the cortex being a network, always operating with a vast number of ongoing changes, i.e. expansions and contractions of states everywhere. Local parts of this network can cross-correlate their expansions and contractions with other local networks in the cortex to form a large set of local networks having a similar expansion and contractions with different lags.

We showed that single trial stimuli and tasks modified the stretching and contractions of the trajectories. These modified fluctuations cross-correlated between particular networks, with the contraction concomitant with the subjective experience. However, our material did not permit a systematic evaluation of relations between task variables, anatomical localization, the expansions and contractions and their spatio-temporal cortical dynamics. Neither was this study designed to examine how local network fluctuations distinguish different single percepts, imageries, thoughts and behaviors.

This dynamic may also be hidden in MEG and EEG signals in human and non-human primates. Future systematic studies should examine the extent to which the modifications of the expansions and contractions and the flow of states predict and distinguish behavioral variables in detail.

## MATERIALS and METHODS

A total of 13 patients (6 females, 7 males; age range: 19 to 55 years; mean: 38 years, SD: 12 years) participated in this study. The patients had drug resistant epilepsy, were candidates for surgical resection and were implanted with depth electrodes, at locations in cerebral cortex determined to reveal the location of an epileptic focus and spread of epileptic discharges. The study was approved by the Human Ethics Committee for the Greater Copenhagen region (permission 56441). The patients spent from 5 to 13 days with the electrodes implanted.

The aim of this study was to obtain field potentials, without any abnormalities, from as many cortical sites as possible. The experimental testing conditions were designed to be as diverse as possible in the clinical ward. The purpose of this diversity was to examine whether the dynamics driving the evolution of the field potential were different for different tests, or explored different parts of state space. The experimental conditions were not designed to map or functionally dissect sensory, motor and cognitive brain functions. The analysis design is summarized in **Figure 1**.

After removal of all epileptic activity (see below), the material comprised 290 electrode leads. The anatomical locations of the electrode leads are listed in **Table 1**.

**Table 1:** anatomical localisation of the electrode leads

| Anatomical site | N<sup>o</sup> leads |
| --- | --- |
| Hippocampus | 38 |
| head (20) |  |
| body & tail (18) |  |
| Middle temporal gyrus | 35 |
| anterior part (8) |  |
| middle part (19) |  |
| posterior part (8) |  |
| Cingulate gyrus | 29 |
| anterior part (19) |  |
| middle part (5) |  |
| posterior part (5) |  |
| Insula | 27 |
| anterior part (13) |  |
| posterior part (14) |  |
| Superior temporal sulcus | 18 |
| anterior part (3) |  |
| middle part (7) |  |
| posterior part (8) |  |
| Superior temporal gyrus | 12 |
| anterior + middle part (10) |  |
| posterior part (2) |  |
| Superior frontal gyrus | 12 |
| Parietal operculum | 11 |
| Temporal pole | 10 |
| Inferior temporal gyrus | 10 |
| Supramarginal gyrus | 9 |
| Fusiform gyrus | 9 |
| Superior frontal sulcus | 9 |
| anterior part (1) |  |
| posterior part (8) |  |
| Parahippocampal gyrus | 8 |
| Orbitofrontal cortex | 7 |
| lateral part (4) |  |
| medial part (3) |  |
| Middle frontal gyrus | 5 |
| Frontal operculum | 5 |
| Lingual gyrus | 4 |
| Entorhinal cortex | 3 |
| Inferior frontal gyrus | 2 |
| Precuneus | 2 |
| Gyrus precentralis | 1 |
| Gyrus postcentralis | 1 |
| Calcarine cortex | 1 |
| Angular gyrus | 1 |
| Subiculum | 1 |

### Experimental Conditions

Typically, 1 to 3 days after the implantation, the patient participated in different tests aimed to induce behavioral brain states as diverse as possible. In addition, we selected episodes of sleep, conversation with the nurse, TV watching, and eating. The purpose of the many behavioral conditions was to let the field potential dynamics explore state space over a wide range. For different practical reasons (medication, post-ictal sleep, early dismissal) not all patients performed all tests. Subjects were sitting or lying in bed. All behavioral conditions were collected several hours away from ictal and post-ictal activity, typically in the first two days in the ward.

#### Visual tests

We collected a series of 200 color pictures of landscapes and outdoor environments, devoid of images of humans (96 dpi, 1060 x 1920 pixels) downloaded from freeware sources on the internet. The pictures were presented on a screen measuring 53×30 cm (refresh rate: 144Hz) which was placed approximately 120 cm from the eyes. Patients were instructed just to watch; with the exception of visual test 6, no behavioral responses were required. A total of 11 visual tests was conducted.

##### Visual test 1

200 pictures appeared, 1 frame each (6.94 ms) with no intermission.

##### Visual test 2

100 pictures were randomly selected and each was presented for 49 ms with no intermission.

##### Visual test 3

the same 100 pictures were presented each for 125 ms with no intermission.

##### Visual test 4

the same 100 pictures were presented each for 125 ms, followed by a 999-ms lasting black screen

##### Visual test 5

the same 100 pictures were presented each for 999 ms, followed by a 999-ms lasting black screen.

##### Visual test 6

The next day, the 100 pictures were mixed with 100 new ones that the patient had not seen before. The 200 pictures were randomly presented each for 999 ms, followed by a 1999 ms lasting black screen. Following each picture, the patient had to indicate whether he/she had seen it earlier (recognition test).

##### Visual test 7

apparent motion. A 3°x 3° black square moved on a white background in steps of 2° from the left to the right border of the screen, in 583 ms. The test was repeated 100 times.

##### Visual test 8

same as visual test 7, but here the square moved from right to left.

##### Visual test 9

one stationary 3°x 3° white square was shown for 1 minute.

##### Visual test 10

one 3°x 3° red square, flanked by four 3°x 3° green squares, was shown for 2 minutes.

##### Visual test 11

one white square, 3 white squares in a row, one red square flanked by 4 green squares, or one green square flanked by 4 red squares were shown for 250 ms. Presentation was random and balanced such that each alternative was shown 50 times. All squares measured 3°x3°.

#### Control conditions

In the Ganzfeld condition, the patients’ eyes were occluded for 10 minutes by white ping pong balls, cut in halves, and covered by surgical gauze pads secured by tape, preventing any patterned visual input. The patients were asked to keep eyes open during the test. In the rest condition, the patient had eyes closed, was encouraged to relax, concentrating on having it black in front of the mind’s eye, but stay awake. Patients were supine in both control conditions.

#### Emotions

A total of 35 pictures (72 dpi) were taken from the Karolinska Directed Emotional Faces (KDEF) test (Lundquist et al.,1998), showing 7 different emotional expressions (angry, disgusted, sad, fearful, happy, surprised and neutral). Each image was shown for 6 s, followed by a 2s dark screen. Following each picture, patients were asked to indicate the type of emotional expression.

#### Memory retrieval tests

During these tests, the patient had eyes closed.

##### Memory test 1

recall faces. With 5 s interval, the experimenter read the names of 29 well-known persons or cartoon characters (e.g. queen Margrethe, Donald Duck, Angela Merkel). The patient responded by yes or no whether he/she could clearly visualize the face.

##### Memory test 2

recall places. With 5 s interval, the experimenter read the names of 15 well-known landmarks or tourist attractions (e.g. the little mermaid, golden gate bridge, the Chinese wall). The patient responded by yes or no whether he/she could clearly visualize the place.

##### Memory test 3

mental navigation. The patient was asked to imagine going out of the front door of his/her residence, going to the right down the street, then walking down the first street appearing to the left and then alternatively to the right and left as in (Roland and Friberg, 1985). After 65 s the patient was asked to tell where she/he is.

##### Memory test 4

same as 3, but starting taking the home street to the left (Roland and Friberg, 1985).

##### Memory test 5

a known target at some distance from the patient’s house was chosen. The patient was asked to imagine walking/cycling slowly to the target and say “now” upon arrival. *Memory tests 6 and 7:* same as memory test 5 but each with a new target.

##### Memory test 8

patient was asked to imagine standing at target (condition 5) and walking/cycling back home. Upon arrival, patient says “now”.

##### Memory test 9

The patient imagines standing at target in condition 6, otherwise as in condition 8.

##### Memory test 10

patient was asked to recall all events in temporal order during the hospital admission day, from getting out of bed in the morning until arriving at the hospital. Upon arrival, patient says “now”.

#### Classification of nouns

A list of 19 Danish nouns all starting with “hoved” (head or main in English) was read (6 s inter-stimulus interval) and the patient was asked to classify each noun as being “concrete” or “abstract”.

#### Motor tests

An EMG disc electrode was fixed to the skin over the forearm extensors, contralateral to the brain hemisphere carrying the largest number of implanted electrodes. A reference electrode was fixed near the wrist, and the EMG signal was sampled by an EMG amplifier (10 Hz-5kHz band-pass filter; amplified and finally recorded at 30 kHz) and used to define the motor epochs.

##### Motor test 1

the patient was asked to move the arm with the electrode such that the hand touches the left side of the chest, to move it back to the original rest position, and then to touch the right side of the chest. Then in similar order, the same hand moves to touch right and left shoulder, right and left foot, right and left knee, right and left abdomen, forehead, right and left heel, right and left thigh, opposite elbow, and opposite hand, in total lasting 60 s.

##### Motor test 2

the patient was asked to quickly bend the index-finger and thumb to touch, then quickly to extend the fingers again and to continue this as quickly as possible for 60s in total.

##### Motor test 3

as in motor test 2, but synchronously with the index and thumbs of both hands.

##### Motor test 4

as in motor test 3, but counterstroke.

##### Motor test 5

rhythm. With the hand contralateral to the hemisphere with the majority of implanted electrodes, the index touches the thumb for approximately one s, followed by 3 short touches and so on in the same rhythm for total of 60 s.

##### Motor test 6

With the same hand as condition 5, the thumb touches shortly the index, middle finger, ring finger and little finger, the little finger, ring finger, middle finger and index and so back and forth for total of 60 s.

#### Attention tests

*Somatosensory attention*: the patient expected a touch stimulus close to detection threshold with a thin calibrated filament (pressure of 10 mg to 20 mg) on the tip of the index finger. The stimulus was first presented after 60 s had elapsed (Roland, 1981).

#### Auditory attention

The patient was asked to listen to a list of random numbers between 1 and 100, pronounced close to his/her auditory threshold. The patient had to repeat each detected number. In the ensuing 60-s period, no numbers were spoken. After 60 s have elapsed, a faint but detectable number was spoken, as a control.

#### Stereognosis

A small object (e.g. a cap of a USB-stick) was placed in the palm of the patient’s hand. The patient had eyes closed and was prompted to manipulate the object with the fingertips. The testing lasted up to 30 s, or until the object was recognized. The test was then repeated with the other hand with another object.

#### Smell and taste

Ortho-nasal smell was tested according to conventional clinical bedside tests using mild odorants such as tea and chocolate. The patient was instructed to sniff deeply, with eyes closed, while the odorant was held close to patient’s nose. Five odorants were used consecutively. Each odorant was tested for 20 s or until the odor was identified, whichever came first.

Retro-nasal smell and taste were tested by the patient pinching the nose with one hand, while he/she placed a sweet in the mouth with the other hand. Mouth-breathing was allowed. The patient was instructed to swallow only saliva, while chewing gently on the sweet. The patient was asked to report the taste of the sweet. After 20 s, the patient was instructed to exhale through the nose, and report any changes in the “taste” (flavor) of the sweet.

#### Sleep

##### 1. Light sleep

Visual confirmation of sleep spindles, 9-16 Hz, minimum duration 0.5 s, maximum 2 s, appearing more frequently than 0.3 Hz in the field potential time series. In addition, visual confirmation of K-complexes > 200 μV, duration >200 ms, diphasic configuration. Finally, the selected 60 s episode should have maximum 20% delta activity. Light sleep was defined as a period of 60 seconds with at least 2 sleep spindles and 1 K-complex. Sleep spindles and K-complexes were detected by an algorithm and afterwards visually confirmed (Andrillon et al., 2011). The criteria for sleep spindles was an increase in 9-16 Hz activity of at least 3 standard deviations, with a minimum duration 0.5 s, and a maximum 2 s.

##### 2. Slow wave sleep

The episode selected had more than 60% slow waves,1-2 Hz > 150 μV peak-to-peak. The amplitudes of the slow waves were more than 3 standard deviations of 0.1-2 Hz activity and had peak-to-valley of at least 150μV occupying more than 20% of the 60 second time period (Hori et al., 2001). All episodes selected were visually confirmed as slow wave sleep.

### Field potentials

Field potentials were sampled at 512 Hz with a band-pass filter of 0.1 to 256 Hz from the implanted electrodes (SEEG electrodes; Ad-tech Medical Instruments, 1.28 mm in diameter, lead length 1.57 mm) in 13 patients. The electrode locations, based exclusively on clinical criteria, were determined from the post-operative CT-scan (1mm^3^ voxels) co-registered to the patient’s T1-weighted magnetic resonance Image (MRI) of the brain. After rejecting electrode leads located in white matter, artifacts and leads with epileptic activity (see below), the field potential from 290 leads located in the cerebral cortex were further analyzed. The sources of the field potential are the transmembrane currents *I*_*n*_(*t*).

From (Lindén et al., 2010):

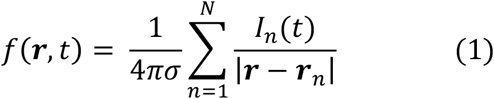

*f*(*t*) is the field potential at a particular instant for a population of neuron membranes close to the electrode lead (Pettersen and Einevoll, 2008). I_n_(t) is the net transmembrane current at a particular distance **r**_n_ from the lead. Under the assumption that the extracellular conductivity, σ, is constant over the whole range of the spatial summation in equation 1, *f*(*t*) at each electrode lead, is the weighted (by their inverse distance from the lead) sum of the membrane and extracellular currents in the sampling space.

The field potential may show net increase of inward currents similar to a net decrease at another electrode, because polarity depends on the reference and might be reversed in recordings. Similarly, inward currents and outward currents in the space sampled by the electrode might cancel, despite lively synaptic activity (Einevoll et al., 2014; Pesaran et al., 2018). Artifacts (synchronous in all leads) and episodes with noise were removed. Thereafter, we used a common reference for each of the gray matter leads. The common reference was the median of the signal in all intra-cranial leads. This efficiently reduces the influence of remote sources on the local field potential. Each field potential then is a time series of fluctuations around an overall median (**Figure1**).

#### Detection and removing inter-ictal epileptic activity from data

To identify epileptic transients in the raw field potential, we performed a wavelet transform on the raw field potential and applied a threshold to the transform to identify epileptic transients. Transient epileptic events were identified using a Mexican hat wavelet (from Matlab^®^ version 2018b). We used three wavelets with half widths of 0.02, 0.04 and 0.1 seconds). These wavelets were then convolved with the field potential signal to generate a time series where the amplitude correlates with transients in data with temporal characteristics of inter-ictal activity. We squared the convolved signal and defined a threshold of 0.5 10^6^ µV as the limit of normal activity. Thereafter all files with detected epileptic activity were removed from the original set of 12949 *f(t)* files at 476 cortical locations, leaving 3476 field potential files from 290 cortical locations each of length 60 s.

### Reconstruction of the dynamics (embedding)

We assume that the dynamics of a local network in the cerebral cortex can be captured from a field potential recording. The dynamics are the set of rules by which the field potential evolves. For a given behavioral condition, the sampled field potential is a time series one can examine in state space to reveal its dynamics. We reconstructed the behavior in state space from the field potentials from the 290 cortical electrode leads for each of the behavioral conditions. In order to find the true dimensionality of the dynamics, we employed the delay embedding method (Takens, 1981). The state of the system at time *t* was represented by a *d*-dimensional vector:

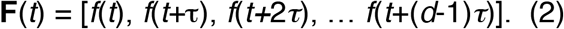

Here, *f(t)* is the field potential. The delay parameter *τ* is taken equal to the empirical autocorrelation time of *f*(*t*) (Pritchard and Duke, 1995). The embedding dimensionality was 20, i.e. d=20, judged enough to capture the underlying dynamics. This method is standard and has a long history in nonlinear system dynamics. The delay-embedded vector signal captures all generic information about the dynamics (Takens, 1981; Pritchard and Duke, 1995). We take this as a working hypothesis here as well.

From each *f(t)*, 20 delayed embedded versions were generated, with delays, τ = 59.9 ms. The delay parameter τ was chosen as the mean of 12949 original field potential time series as the value where the autocorrelation function of these time series = 1/e (Pritchard and Duke, 1995) (**Fig S1**). For each behavioral condition, the first 60 s of the *f*(*t)* was chosen as the basis of the embeddings.

### Dimensionality of state space

The dimensionality of the dynamics was estimated with the Grassberger and Procaccia (1983) method by an automatic algorithm developed in-house. Specifically, for each point of the embedded time series, the distances to all other points in the state space were calculated. The number of points within a radius of ε was then counted for different values of ε. The correlation integral is given as:

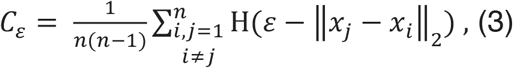

where *n* is the number of points, *x*_*i*_ and *x*_*j*_ are *d*-dimensional coordinate vectors, *ε* is the radius threshold and *H* is the Heaviside step function. The || indicates the norm. The calculation of *C*_4_ is computationally intensive but parallelizing the code made it possible to reduce the computation time using the Danish Super Computer center Computerome (http://www.computerome.dtu.dk). The algorithm arrives at the state space dimension by considering the dependence between the correlation integral and the distance thresholds for a given embedding dimension. Specifically, the slope of each curve produced by a log(C_E_) and log(ε) plot is automatically examined to determine the correlation dimension of the upper linear part of the curve **(Figure S1)**. The automated program for this identification has a manual counterpart since, in a small number of cases, it was difficult to determine the slope automatically. Finally, to determine the dimension, the dependence between the correlation dimension and embedding dimension is considered. If the correlation dimension is constant for 3 or more successive embedding dimensions, the algorithm concludes that the associated correlation dimension is the state space dimension, in short correlation dimensionality (**Figure S1)**. The constant part of this dependence is determined by a least squares approach (**Figure S1**). Both the automatic detection of the linear slope and the automated horizontal line determination of the dimension have multiple tuning parameters that can be more or less restrictive depending on the amount of noise in the data. For 28 records it was not possible to find a horizontal segment.

The correlation integral estimate depends on the number of sample points, here 30720 (60 s). Increasing the number of samples will increase the dimensionality slightly (**Figure S1**), but further decreases the likelihood that the dynamics, and hence the dimensionality, remains (quasi) stationary. So, the 60 s is a compromise.

The correlation integral algorithm also determined the amount of false nearest neighbors, and the inequality R_d+1_(n)/R_A_ > A_tol_ (see Kennel et al., 1992 for all details). If the embedding dimension chosen, in our case 20, is too low, the trajectories will intersect and overlap in state space. Such intersections and overlaps are called false nearest neighbors. False nearest neighbors thus tell that the embedding dimension chosen to reveal the dynamics is too low. R_A_ is an estimate of the size of the set containing all trajectories, R_d+1_(n) the sum of distances between each of the n points and neighbors, and the A_tol_ set to = 1 (Kennel et al., 1992). When the number of dimensions increase the distances between the points in state space increase; this is especially true for time series with a limited number of samples. The above inequality limits the possibility that false neighbors at high dimensionality become true neighbors. At embedding dimension 4 the number of false nearest neighbors was practically zero and the A_tol_ criterion fulfilled up to embedding dimension 20 (**Figure S1**). This means that the **F**(t) unfolds as expected from a dynamical system at embedding dimension 4 such that there are no crossings (intersections) of the trajectories in the attracting object (Kennel et al., 1992). For comparison, we shuffled the original time series to produced surrogate data with the random method (Matlab® randperm) (**Figure S1**).

The correlation dimensionality estimate seems superior to the alternatives (Camastra, 2003). The extraction of the dynamics from the embedded times series of the field potential was done by using integer dimensionality close to the estimated correlation dimensionality. With the absence of false nearest neighbors and the A_tol_ criterion fulfilled, these factors together should give a close to exhaustive description of the dynamics contained in the field potential.

### Statistical analysis of dimensionality

When the distribution of estimated correlation dimensions over all behavioral conditions was plotted for each electrode lead, it appeared that in some cortical locations, the mean correlation dimension was high and the square-root of the correlation dimension variance relatively low; whereas in other locations mean correlation dimension was lower and the square-root of the variance relatively high (**Figures 2A, S2**). For each patient, we checked whether the distributions of the mean correlation dimension and square-root of the variance were approximately Gaussian. Under the assumption that the correlation dimensions distributed in two groups, one with high mean and low standard deviation, standard deviation, and one with lower means and higher standard deviation we subjected the correlation dimensions for each patient separately to a k-means cluster analysis (k=2). Two different centers were selected and a distance from the center to each point is measured. The algorithm then aims to have the lowest possible sum of distances. The algorithm is given as:

1. Initialize: select *k* points as medoids
2. Compute distances of each data point to each medoid
3. Assign each observation to the cluster with the closest medoid.
4. Compute average of the observations in each cluster to get a new medoid location
5. Repeat step 2-4 until cluster centers does not change

The medoid method implies that the algorithm finds one experimental co-ordinate, with minimum distances to all members of that cluster, as the center point for each cluster (which reduces the influences of outliers). **Figure S2** shows the results. The quality of the clustering is apparent from the silhouette plots (Hastie et al., 2009). The silhouette value ranging from –1 to 1 shows how similar a cluster point is to its own cluster. A value a bit above 0 is usual in accepting the clustering. Silhouette values are shown in **Figure S3**.

To further statistically characterize the separation of the two groups we made a linear regression of the points of each cluster and compared the regressions lines (F-test) and verified that the residuals were approximately distributed as a Gaussian distribution. **Table S1** shows the results.

### Dynamics confined to one part of state space

**Figure 3A** shows all 106,782,720 normal states in one state space. These states were distributed in 3476 sets. As the set of points occupying state space were multidimensional, their occupancy of state space was approximated. If the dimensionality was 6.6, for example, the set of points were located to a 6-dimensional state space, because the states do not exist in dimension 7. Each of the 3476 sets including all states was covered in a convex hull, the smallest convex set that contains all 106,782,720 states. The convex hull was computed for all relevant embedding dimensions. This was done using the Matlab^®^ function convhulln using tetrahedra forms as facets. For the illustration, each of the 3476 sets was assigned one color; dimensions 1 to 3 and 4 to 6 were plotted separately in the figure (**Figure 3A**).

### Fluctuating dynamics

With data sampled over 60s, the sets plotted in 3,4,5,6, or 7 dimensions will be unable to reveal fast dynamics relevant for perception, thinking, and preparation and execution of behavior. To examine faster changes in state space occupancy, we approximated location of the states as a hyper-ellipsoid (see **Figure 3A**). Mahalanobis distance is a measure of distances between data points in a multidimensional space, that is scale invariant and independent of Gaussian assumptions. Each state has a Mahalanobis distance in a multidimensional space from the center of the hyper-ellipsoid of the integer dimension just lower than the correlation dimensionality. We calculated the enclosing hyper-volume, of each of the 3476 sets.

We also used the *embedded time series* **F**(t) in the appropriate integer dimension to estimate the volume of the attractor (attracting set of states). If the correlation dimensionality of the set was 5.8, we calculated the first 5 principal components of the 5.8 dimensionality set. In this case, the estimated hyper-volume of the set in 5-dimensional state space was calculated as:

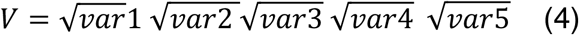

in which the product of the square-roots of the variances in each of the 5 orthogonal principal components dimensions is a hyper-rectangle giving the estimate V of the set in state space. The algorithm determined the V every 50 ms with a certain sliding window (was set to 128 samples = 250ms). The 50 ms are less than τ (60 ms). The sliding window was less than 5 τ, because most data ranged between 5 and 7 dimensions (see Flow of states below). This means that each value of V is an estimate of the volume of 25 consecutive states. Note that this method estimates V as the volume of an orthotope (hyper-rectangle) and not of a hyper-ellipsoid. In equation 4, the orthotope is 5 dimensional. In addition, the succession of the 128 states was confined to a trajectory, that however could take various shapes (as also seen for the 1 s and 5 s trajectories in **Figure 3**). The states, however over some 10s to 20 s become well bounded by a hyper-ellipsoid as the convex hull. V fluctuated roughly from 10^7^ to 10^17^ μV.

The volume expansions and contractions of local cortical states in state space were estimated in successive 50 ms intervals. Since we used only 25 states for the calculation of the volume, it made no sense to narrow the observation intervals. This does not preclude studies of faster fluctuations of the expansions and contractions, if the sampling rate of the data is sufficiently high and if the autocorrelation in the time series is rapidly declining. The volume of the states varied significantly with the dimension of state space. Most likely, the dimensionality of the dynamics changes faster than over 60 s, the time needed to get a reasonably accurate estimate. However, the volume of the cortical states may reflect smaller changes in dimensionality. If the distances between states increase, the product of the square roots of variances and hence the volume of the states increases (equation 4). This is roughly equivalent with an increase in dimensionality. Similarly, if the distances between states decrease, the variances and hence the volume of the states decrease. With our data we could only observe volume fluctuations slower than the average flow of states. However, the flow of the states did not seem to fluctuate faster than the volume (see **Figure S6** values).

The amplitude variations of the fluctuations of the log _10_ V from the median in all 60 s test episodes in all awake states (n= 3526) in all patients (n=13) were measured and shown in **Figure 3D**. For comparison, the similar fluctuations from 20 randomly selected 60 s episodes in awake states, i.e. spontaneous behavior not belonging to any test episode, were also calculated (**Figure 3D)**. A coefficient of variation of the z-score values of V (see below) was also calculated in steps of 50 ms with a sliding window of 250 ms (**Figure S5**).

### Attractor properties

The set of states as points occupying state space were located to one part of state space. Neither the states themselves nor the trajectories they formed did spread to parts of state space outside the convex hull in **Figure 3A**. The reason for this could be that the network in which the states evolved had attracting properties, i.e. attracted states nearby. If we defined the hyper-ellipsoid as being within 90% limits of the Mahalanobis distances, the 10% external states (outliers) should sooner or later end up in this hyper-ellipsoid if the local network had attracting forces. We used the Matlab^®^ function mahal. The Mahal_Threshold_ = 90% percentile of Mahalanobis distances of all states from the center of the hyper-ellipsoid. The states included in the hyper-ellipsoid < Mahal_Treshold._

**Figure S4G** shows the probability densities of all states. We defined the putative attractor as the enclosed volume V_90%_ containing 90% of all Mahalanobis distances. Thus 10% by definition were outliers.

Since the hyper-volume, V, itself fluctuated over a scale from 0.2 to 10 Hz, during which V ranged from small to maximal, the attracting property should be universal, present when V was small, medium and large. As the distribution of fluctuations in log_10_V was near Gaussian (**Figure 3D**) we defined small logV_90%_ as the lower 16 % of the distribution of the 90% Mahalanobis distances in each patient, medium as the mid 68% logV_90%_, and large as the top 16% of logV_90%_. The algorithm then located small medium and big states of logV_90%_ and listed the length of the outlying states, i.e. the time in ms from departure to the return to the hyper-ellipsoid. This was done for all conditions and electrode leads (n=3476) **(Figure 3E, F, G)**.

*Flow of states*. The **F**(t) was visualized in the integer dimension just lower than the correlation dimensionality. If the correlation dimensionality was 6.6, for example, we chose 6 dimensions for visualization (**Figure 3**). Six dimensions gives 6 orthogonal planes (**Figure S4**). The planes were defined by their normal vector and origin. The set contains only one, in this case, 6-dimensional trajectory, describing the evolution of states. Note that each state is identified in state space as, in this example, a 6-dimensional coordinate [μV_*ti*_, μV_*ti*+1_τ,μV_*ti+*2_*τ*, μV_*ti+*3_*τ*, μV_*ti+*4_*τ*, μV_*ti+*5_*τ*]. The states in state space progress every 1/512 s from one 6-dimensional μV value to the next. The 6-dimensional μV value however is composed from 6 samples spread in time over 360 ms. This is how the flow of states must be interpreted. If the Euclidian distances between successive 6-dimensional μV values increase, the flow of states increases. When Euclidian distances between successive 6-dimensional μV values decrease, the flow of states decreases.

A relatively detailed description of how the trajectories behave can be obtained from calculations of how long time it takes for states to cross one of the symmetry planes (**Figure S4**). As an example, depending on where in this 6-dimensional space a continuous set of states are, they may first cross for example the 3^rd^ symmetry plane (from one side or the other), after a while they may cross the 1^st^ symmetry plane and after yet a while the 5^th^ and so on. The cross time is the number of states positioned on one side of the plane until they cross that plane (**Figure S4C, E, Figure S6**). Cross times were calculated for the whole material of 60 s files (n=3476) (**Figure S6)**.

To further prove the presence of attracting forces, we calculated flow vectors (Forsberg et al., 2016) as the average velocity over 4 samples (7.81 ms). In **Figure S4**, only the flow vectors after the time point where the Mahalanobis distance was maximal are shown. We calculated also the Euclidian distance in μV between the point, where they exceeded 95% limit and the point of return to the enclosing volume in state space for all 3476 files (**Figure S4**).

### Single trial analysis of the fluctuating attractor

For the conditions in which the patients were stimulated by visual stimuli (visual tests 4, 5, 6, 11; emotions) the start of the stimulus is shown in **Figure 6**. For the conditions in which the patients were verbally prompted to recall faces and places (memory retrieval tests 1, 2) and the classification of nouns, the trials are marked at the peak in the analog sound track (sampled with 30 KHz). For the visual stimuli conditions, the single trials were aligned to the start of the stimulus from the photodiode analog track (sampled with 30 KHz). An epoch from -4000 ms to + 4000 ms was chosen. For each of these epochs, the log V signal was converted to z-scores by the Matlab^®^ function z-score. Using the peak in the sound track (providing the most well-defined trigger signal) to align all single trials in the 60 s episodes, segments, 8000 ms long, of z-scored V were calculated. Thereafter all 8000 ms segments, one for each trial, were averaged and the average V plotted (**Figures 6 and 7)**. For the conditions in which the patients were verbally prompted to recall faces and places (memory retrieval tests 1,2) and the classification of nouns, the single trials were aligned to the peak in the sound track and similarly treated. This was done for each lead.

For each behavioral condition, the cross correlation between all leads of their individual average z-score was calculated by the Matlab^®^ function xcorr using the option coef, for the time interval -1000 ms to + 2000 ms. Thereafter the pairwise lag between any 2 leads was calculated. A 95% confidence interval of the pairwise cross correlations served to separate significant pairs from random fluctuations.

### Anatomical Registration

Image registration was applied for each patient’s images in order to compare all the subjects in a unique reference space. The applied procedure is depicted in **Figure S7**. The MNI152 atlas is used as the global reference standard brain (Fonov et al., 2009). Each individual MRI image with all electrode leads marked was transformed into standard format and merged with the MRIs in standard format of the other patients.

## Acknowledgments

We gratefully acknowledge the expert technical help of Lennart Derm and Thomas Kirkeskov Olsson, Dept. Clinical Neurophysiology University Hospital Copenhagen and the editorial help of Paul Cumming. We are greatly indebted to John Hertz for his thoughtful discussions and important input on earlier versions of the manuscript, to Aasa Feragen for advice on multidimensional volumes, to Laurence Dricot for help with the anatomical localization of the electrode leads, and to Viktor Jirsa for his critical reading and comments.

## Funding

This study was supported by the Velux Foundation, Denmark.

## Author contributions

PER conceived and designed the study. PER and AS got the funding. BJ did surgery implantation and postoperative survey. MF was responsible for field potential acquisition and wrote the ethics application with AS and PER. AW created all data analysis pipelines and wrote all software together with SNL SH, CKKH. JM and AW developed hardware. Data acquisition: JM, AW, CKKH. Experimental testing of patients: JM, RK, PER. PER, AW, SNL developed the attractor volume model. AW, PER, JM, CKKH did the data analysis. SNL did statistics. RK, JTD did the anatomical analysis. RK made all Figures. ML, LP, AS managed and controlled the clinical conditions of the patients. PER wrote the earlier versions. PER, JM and RK edited the final version of the manuscript.

## Competing interests

Authors declare no competing interests.

## Data and materials availability

data are available at **github** https://github.com/ChristofferKKHansen/RolandLab.

## Supplementary Materials

Figures S1-S7

Tables S1-S2

## Notes

https://github.com/ChristofferKKHansen/RolandLab

### Competing Interest Statement

The authors have declared no competing interest.

