## Supplemental figures and Tables for "Human cerebral cortex networks use expanding and contracting state dynamic to shape cortical functions"

**This PDF file includes:**

Figures. S1 to S7
Tables S1 and S2

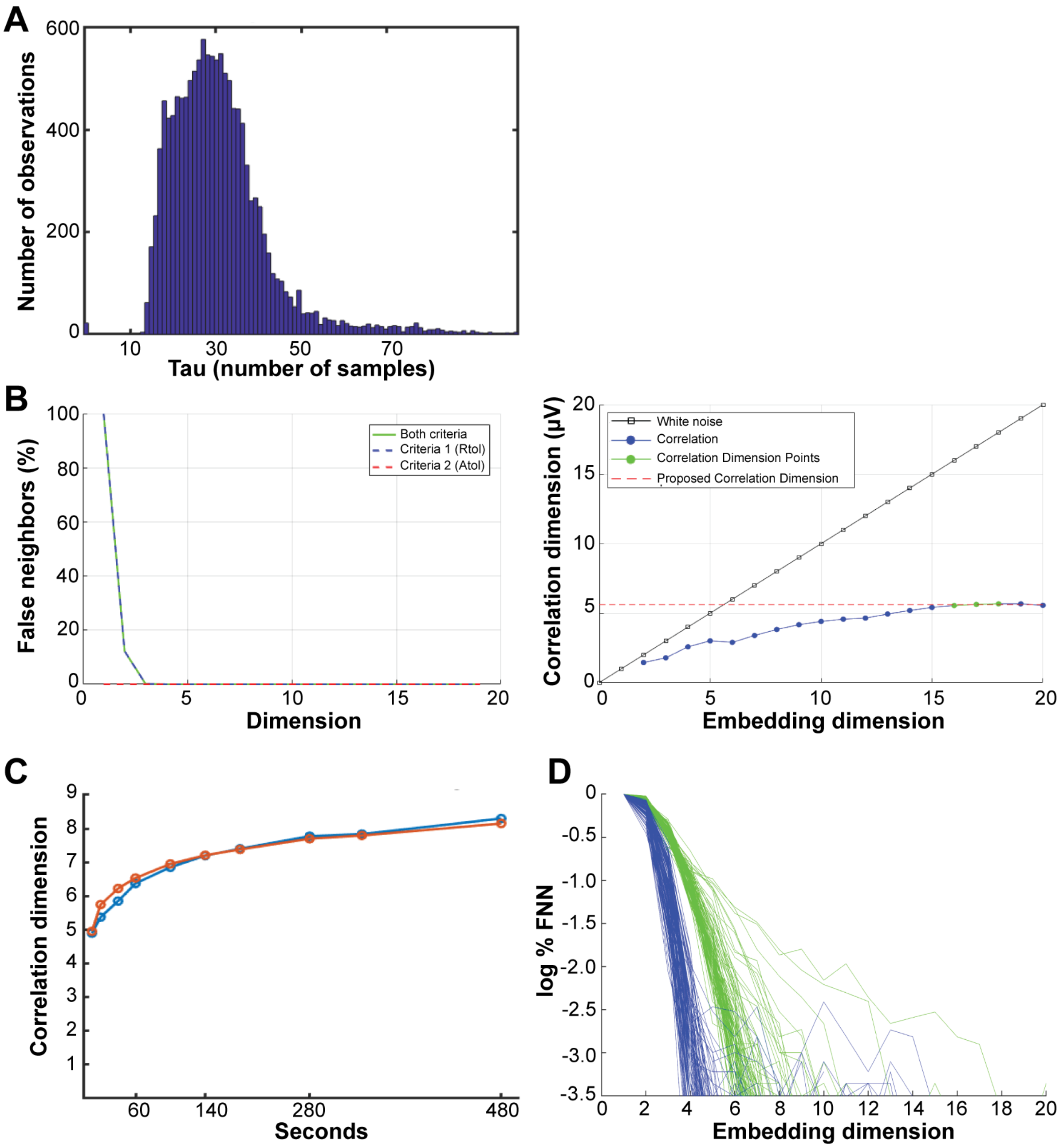


**Fig. S1. Estimation of the dimensionality of *f*(t). A.** Distribution of 12949 optimal values, chosen at the time when the autocorrelation function equals 1/e in the original f(t) time series. The $\tau$ mean = 30.65 samples; equivalent to 59.9 ms. All correlation integrals were calculated with $\tau = 30$ samples. **B.** Algorithm for estimating the correlation dimension of the state space. Left: false nearest neighbors and the $R_{d+1}(n)/R_A$ criterion as function of number of embedding dimensions. Right: the algorithm finds the first horizontal segment of the curve relating embedding dimension to correlation dimension. **C.** Estimated correlation dimension as a function of data samples. Data from patient 4, lead 1 (blue) and lead 53 (red). With increasing number of samples, the correlation dimension continues to increase, presumably a consequence of decreasing stationarity. **D.** False nearest neighbors (FNN) in original (blue) and shuffled data (green).

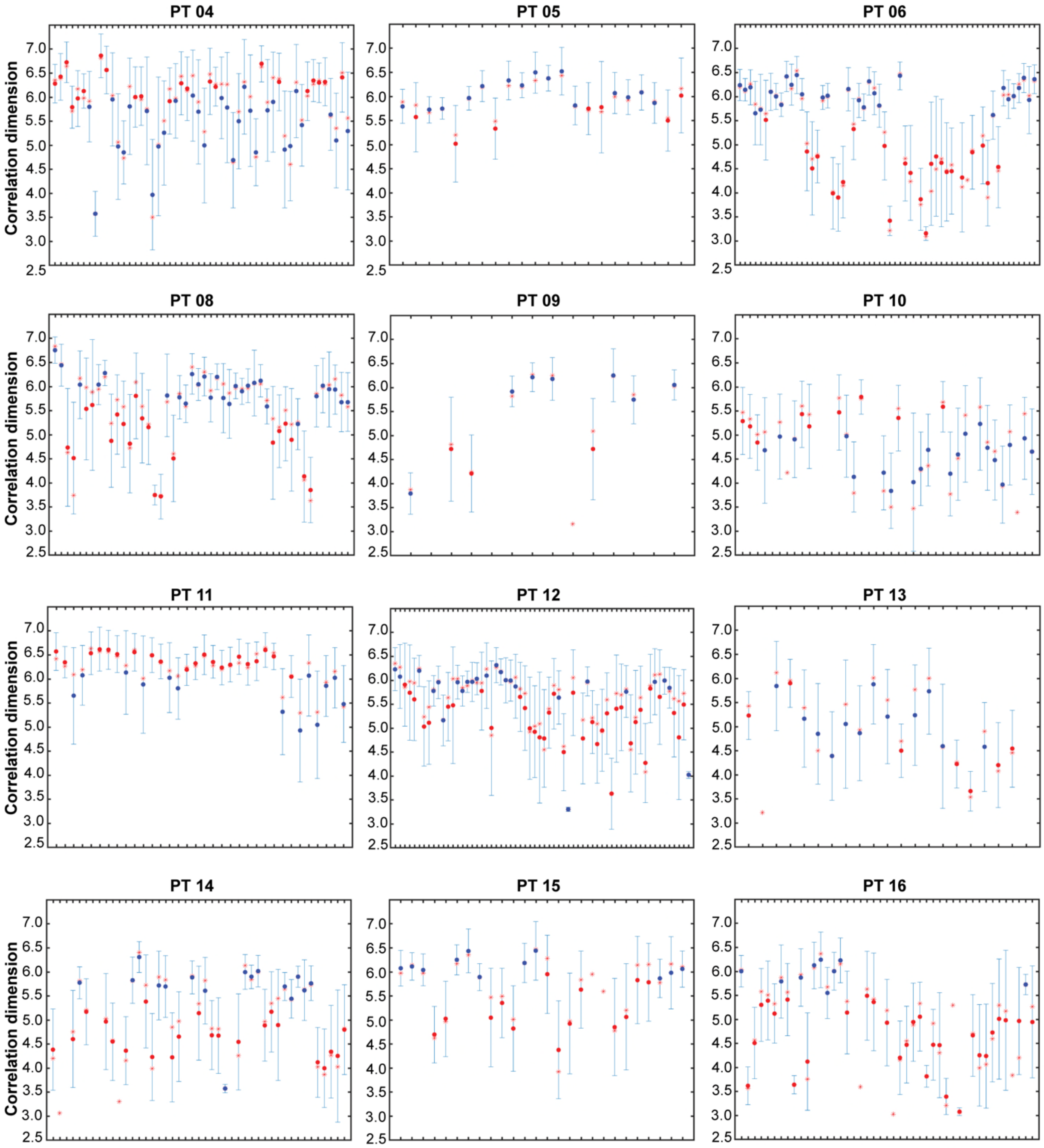


**Fig. S2. Distributions of the mean correlation dimensions**. Correlation dimensions for all tests completed by the patient along the y-axes; x-axes: positions of electrode leads (not grouped anatomically). Red star: median. Red or blue dots: mean. The dot color depends on the membership of cluster (**Figure S3**). Standard deviation shown as error bars.

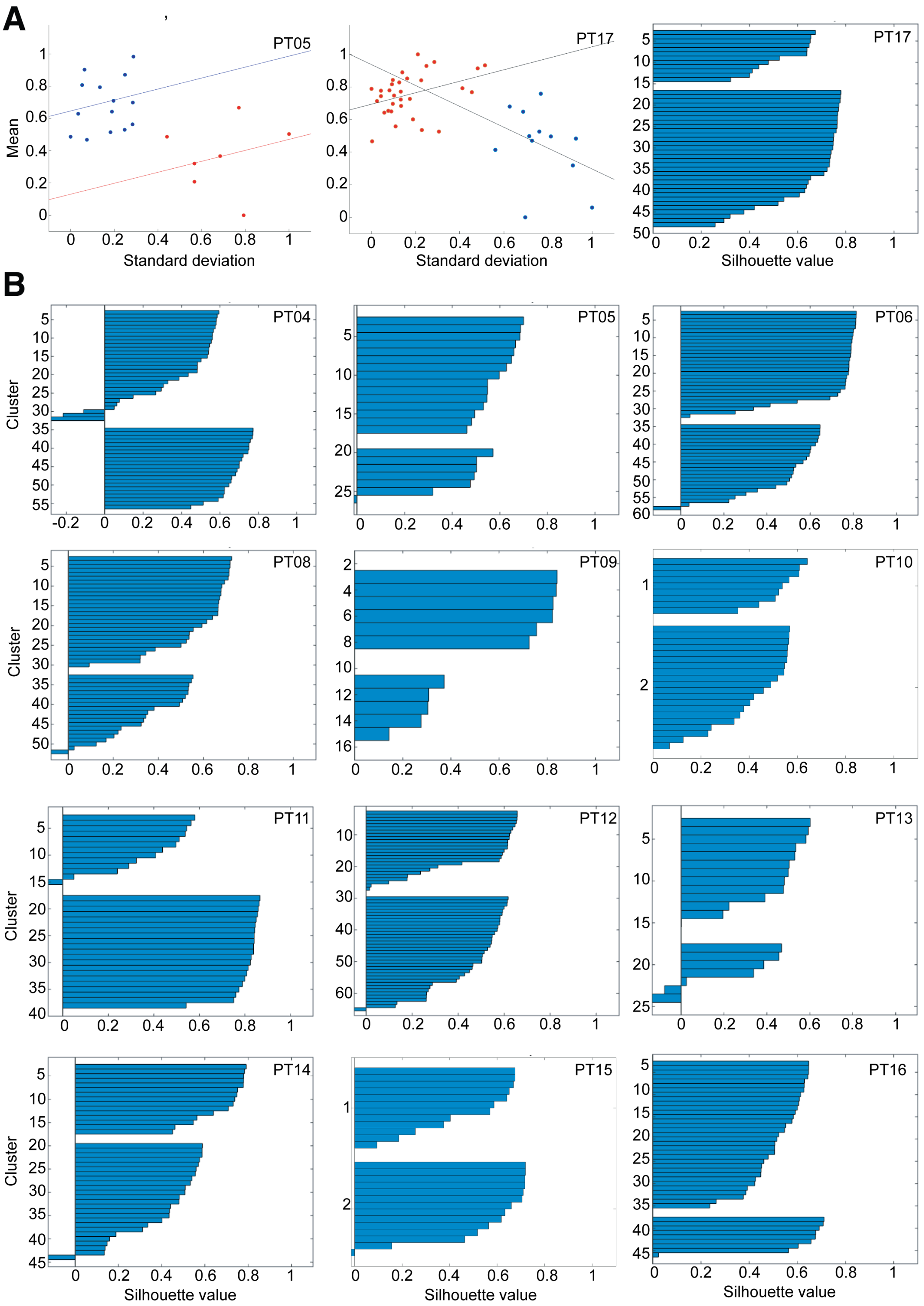


**Fig. S3. Two distributions of correlation dimensions: high mean, small standard deviation (SD) and lower mean and larger SD. A.** Left. Blue dots: high means, low SD, with regression line; red dots: low mean, high SD, with regression line. Middle. Red dots: higher mean, lower SD; blue dots: lower mean and higher SD, with respective regression lines (**STAR Methods**). Right: silhouette plot of the clustering in A middle. **B.** Silhouette plots for the medoids k-2 clustering into 2 groups in 12 patients.

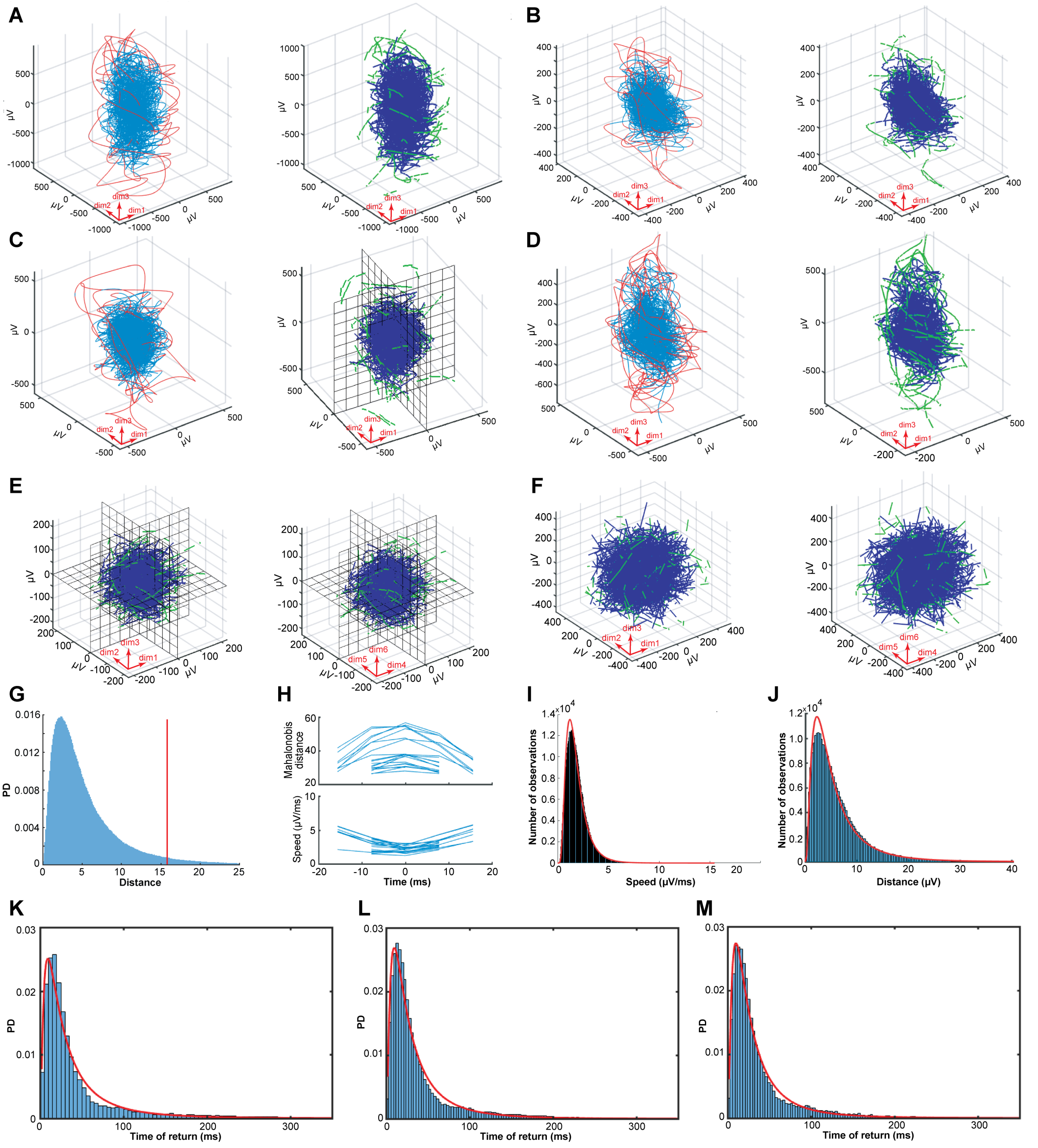


**Fig. S4. Attractor trajectories, flow and properties.** In **A** to **D,** the fluctuating attractor is defined as the hyper-ellipsoid containing 95% of all states over 60 s. When the trajectory is inside the attractor, it is blue, when it is outside it is red. Only projections of the first 3 dimensions are shown. Panels are from 4 different patients. In A to D, attractor trajectory and vector flows (**STAR Methods**) are shown to the left and right, respectively**.** Vectors inside the attractor are blue and vectors outside pointing towards the attractor are green. **A.** Slow wave sleep, Correlation dimensionality (CD): 2.85, anterior part of insula. **B.** Recall places, CD: 5.85, anterior part of cingulate gyrus. **C**. Motor sequence (motor test 6), CD: 6.05, mesial part and anterior part of superior frontal gyrus. Right: flow vectors with 2 symmetry planes (different angle of view chosen for clarity). **D.** Eating, CD: 5.81, amygdala. **E.** Flow vectors pointing towards fluctuating attractor in projections of 6 embedding dimensions with corresponding attractor symmetry planes (as used in **Figure S6** for the calculation of flow of states inside the attractor). Judging the moods from facial expressions, lead 24, cortex lining posterior part of superior temporal sulcus. Left: first 3 dimensions; right: dimensions 4,5,6. **F.** Flow vectors, first 60 s of mental navigation (test 8), lead 25, frontal eye field. **G.** Histogram of Mahalanobis distances from the center of the hyper-ellipsoids for all states in all 3476 combinations of leads-conditions with normal field potentials; y-axis: probability density (PD). **H.** Representative distance and velocity profiles of departures from the fluctuating attractor, aligned to maximal distance from the attractor (time = 0). lead 4, imagery of faces. **I.** Logarithmic-normal distribution (red curve) of speed of all flow vectors (n = 506,409) towards all attracting sets. **J.** Log-normal distribution of distances (n = 253,490) between the point of departure and re-entry into the fluctuating attractor (**STAR Methods**). **K.** Log-normal probability density (red curve) of times to return to the attractor for 10% outlying states when the attractor V is small; median 23 ms. **L.** Idem for medium size V, median 24 ms. **M.** Idem for large V, median 23 ms.

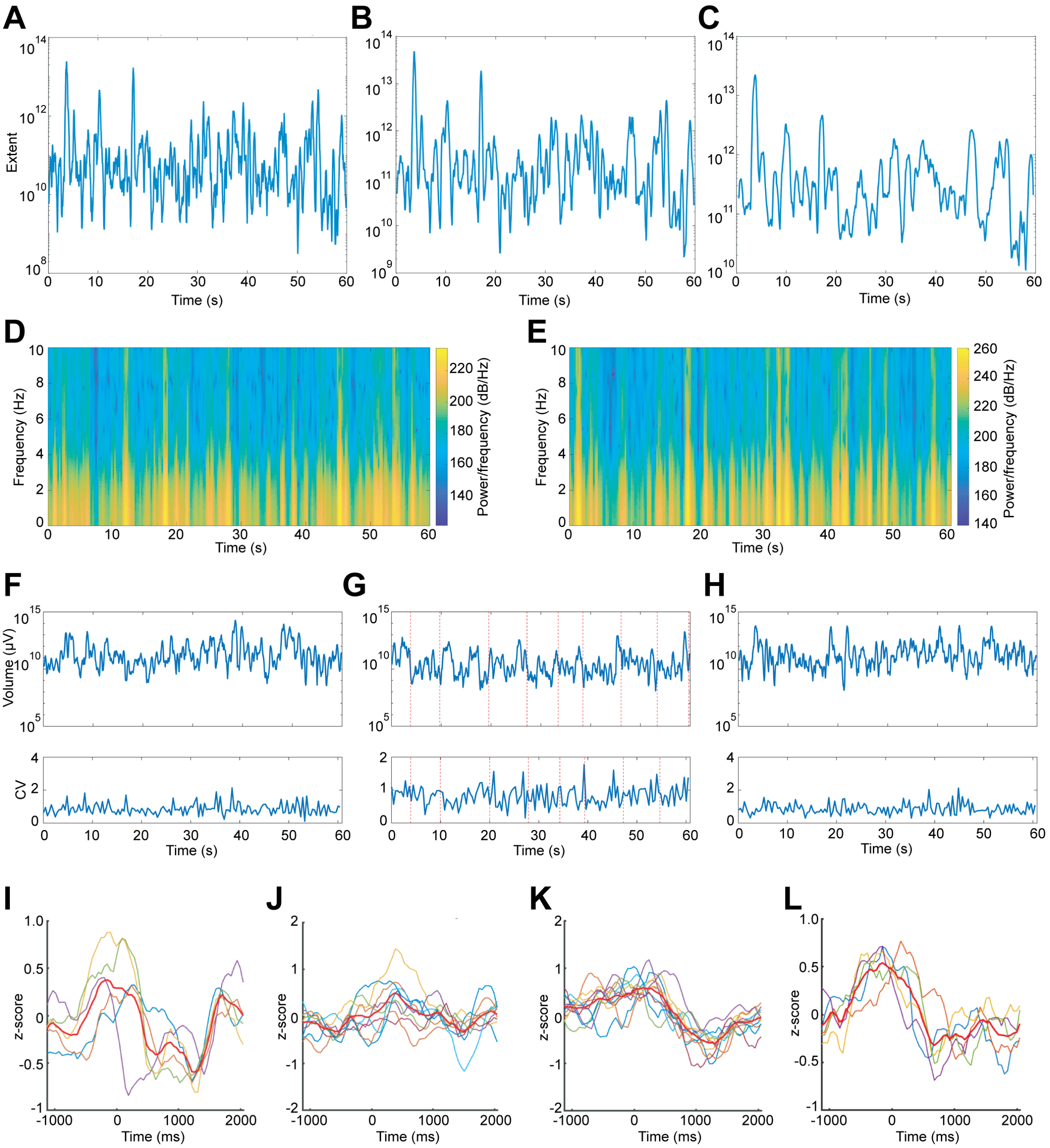


**Fig. S5. Expansions and contractions of the volume of the fluctuating attractor.** Panels **A** to **H** show the volume (V) calculated according to equation 4 (STAR Methods) to show fast fluctuations of the attractor volumes**.** V is thus calculated as the volume of 25 consecutive cortical states. **A.** The evolution of V with a sliding window width of 128 states. **B.** Sliding window width of 256 states. **C.** Sliding window width of 512 states. **D.** Power spectrum of fluctuations of expansions and contractions of the attractor, classification of nouns (patient 12, lead 24), supramarginal gyrus. **E.** Power spectrum, judging the mood of faces (patient 17, lead 2), amygdala. **F.** V, and for comparison, the coefficient of variance of the z-score of $\log_{10}V$, CV (patient 8) rest, anterior hippocampus. **G.** Same patient, same electrode, classification of nouns, red vertical lines mark the peaks of the sounds of the nouns. **H.** Same patient, mental navigation (test 4), fusiform gyrus. F, G, H show typical ranges of the CV of $\log_{10}V$ between 0.5

and 2.5. However, the irregularity of V itself is orders of magnitude larger. **I.** Cross-correlated leads, z-scores (patient 14) recalling places. Different leads have different colors. The mean of the average z-scores is shown in red. **J.** Non-significant leads (patient 14) recalling places. **K.** Cross-correlated leads (patient 5) judging the moods of faces. **L.** Cross-correlated leads (patient 12) classifying nouns.

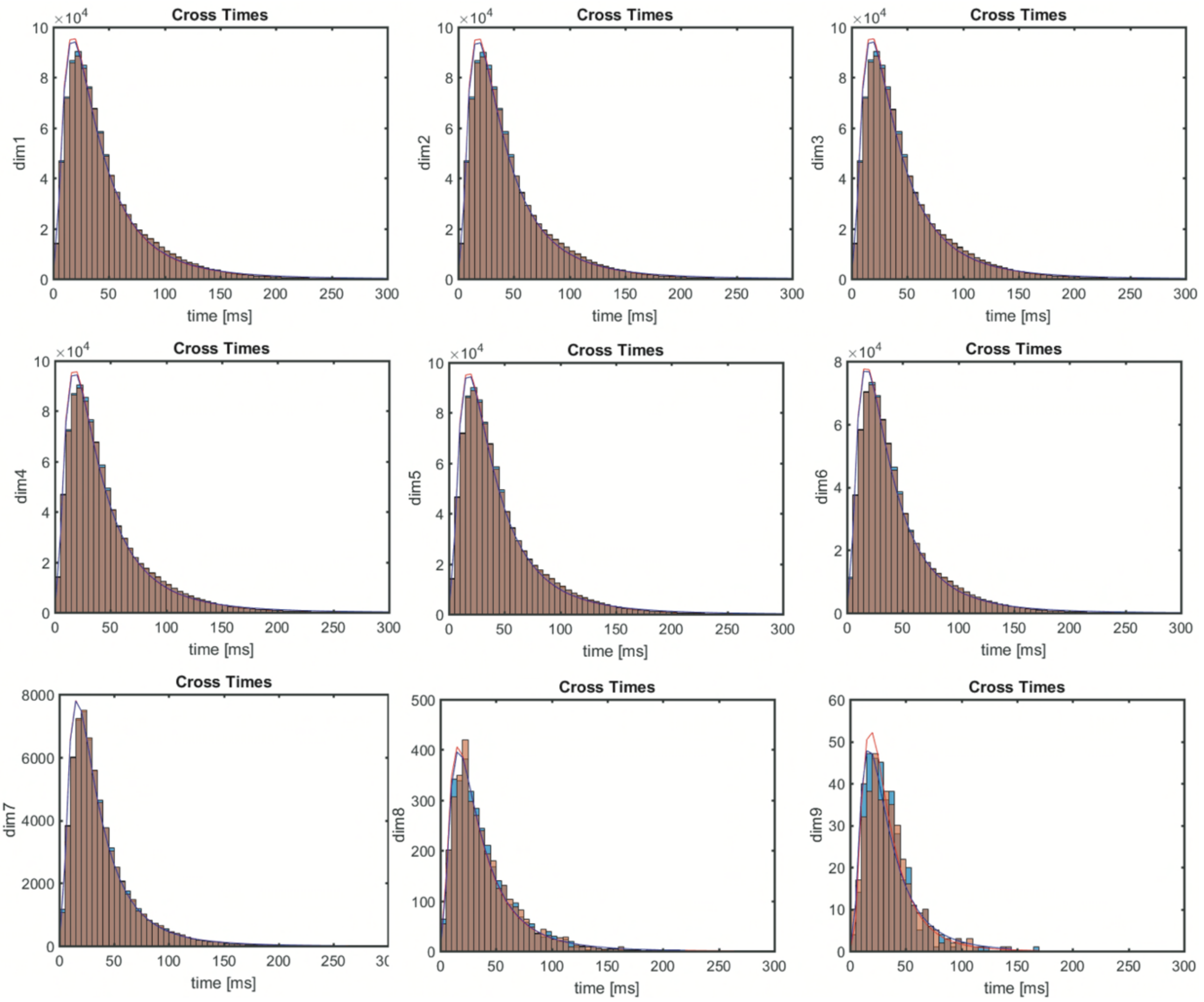


**Fig. S6. Flow of all states in the attractors with respect to symmetry planes.** The flow of cortical states inside the attractor reveals the details of the dynamic. The panels show the distributions of the time–states spend in one sector until they cross the symmetry plane and continue into a new sector of the attractor, for all 3476 attractors. **Figure S4 C** and **E** show examples of attractors with symmetry planes. Light brown: flow from the positive side to the negative; blue: states flowing from the negative to the positive side; brown overlap of the two distributions. All distributions were logarithmic normal (continuous curve: best fit). Median values for dimension 1 to 6 were 34 or 35 ms, for dimensions 7,8,9, medians were 30 and 31 ms. Together with **Figure 3,** this limits the possibilities of alternative interpretations of the dynamics of the data (for example fixed point attractors, bifurcations, limit cycles, tori, multi-stabilities and longer lasting trajectory condensations).

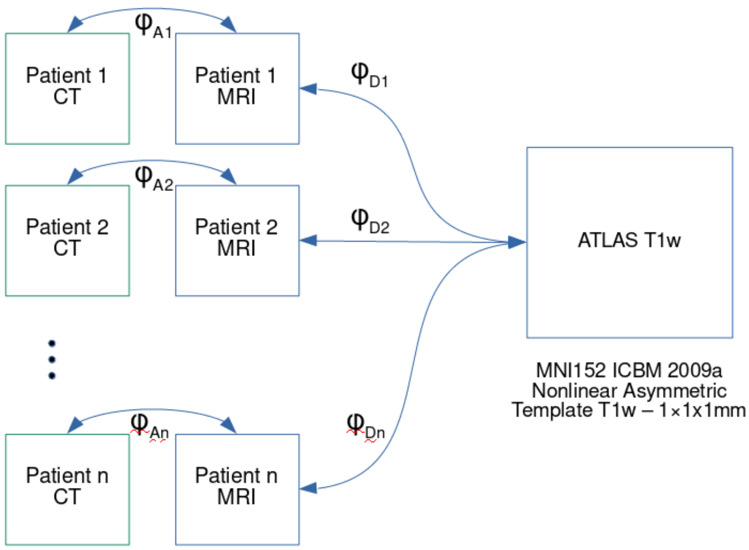


**Fig. S7. Image registration pipeline using MNI152 ICBM 2009.** The template used was the nonlinear asymmetric T1-weighted ICBM 2009, with a resolution of 1x1x1mm (Fonov et al., 2009). The CT and MR images were registered with an affine transformation ϕAi, where i = 1, ..., n refers to the patients. The affine registration algorithm is intensity-based and implemented with ITK (ITK, 2019). The similarity metric is mutual information (Thévenaz and Unser, 2000). The algorithm computes a rigid transformation first, and then the affine transformation in a multi-resolution framework. The affine transformation was applied to the CT images to align them to the MR images. Prior to atlas registration, the MRI images were segmented. Brain extraction was performed using the ROBEX algorithm (Iglesias et al., 2011). The MRI images were registered to the atlas with a deformable transformation, ϕDi. The deformable image registration method used was Symmetric Image Normalization (SyN) (Avants et al., 2008) which uses cross-correlation (CC) as similarity metric and the L2 norm of the velocity field as regularization. The transformation or displacement field ϕDi was computed by integration. Finally, the deformable transformation was applied to the CT and MR of each image patient.

**Table S1. P-values, F-test difference clusters**

| | |
|---|---|
| PT04 | 0.00026792 |
| PT05 | 0.0099752 |
| PT06 | 2.6235e-14 |
| PT08 | 8.2901e-09 |
| PT09 | 1.3745e-07 |
| PT10 | 1.4378e-08 |
| PT11 | 0.56562 |
| PT12 | 0.0018379 |
| PT13 | 0.020649 |
| PT14 | 3.8273e-13 |
| PT15 | 0.00024565 |
| PT16 | 1.2043e-09 |
| PT17 | 0.0042629 |

Difference between the linear regressions of the  dimensionalities forming the two clusters (examples in **Figure S3** top)

### Table S2. Single trial tasks

| Patient | Test | Active electrode leads |
|---|---|---|
| 4 | visual test 4 | 1,4,10,15,16,18, 21,22,23,29,33,37,40,49,53,54,57,65,68 |
| | visual test 5 | 4,18,21,22,29,37,409,57,68 |
| | visual test 6 | 1,4,10,15,16,18,21,22,28,29,33,49,57.65,68 |
| | visual test 11 | 4,10,15,18,20,28,37,40,49,57,65,68 |
| | emotions | 49,53,54 |
| | recall faces | 1,22,37,40,57,68 |
| | recall places | 21,23,40,49,57 |
| 5 | visual test 4 | 1,2,5,6,7,10,11,14,18,19,24,25,26,39,40,49,54,55,56,57,59,65 |
| | visual test 5 | 1,10,11,14,18,19,24,25,26,54,55,56,57,65 |
| | visual test 6 | 7,10,11,14,18,19,24,25,26,39,40,49,55,56,57,59,65 |
| | visual test 11 | 1,25,26,39,40,54,65 |
| | emotions | 1,2,5,6,10,11,24,25,55,56,65 |
| | recall faces | 1,5,6,7,11,14,18,24,26,39,55,56,65 |
| | recall places | 1,18,26,54,65 |
| 6 | visual test 4 | 1,2,3,15,17,51,57,105,115,116,117 |
| | visual test 5 | 1,2,3,8,15,17,26,51,57,105,115,117 |
| | visual test 6 | 1,2,3,15,51,57,58,105,107 |
| | emotions | 67,105,107,115 |
| | recall faces | 1,2,3,8,15,2651,57,105,115 |
| 8 | visual test 5 | 9,26 |
| | visual test 6 | 69,70,79 |
| | recall faces | 2,20 |
| | recall places | 73,96,97,98,99 |
| | nouns | 66,69,70,74,79 |
| 9 | recall faces | 21,61 |
| | nouns | 21,22,61 |
| 10 | recall places | 39,42 |
| | nouns | 12,39 |
| 11 | visual test 4 | 36,41,46,47,48,50,53,56,68,69,88,101 |
| | visual test 5 | 1,2,9,21,23,24,33,34,46,47,48,49,50,53,56,68,69,88 |
| | visual test 6 | 8,10,21,23,24,25,41,46,47,48,49,50,53,68,69,88,93,101,102 |
| | visual test 11 | 2,8,9,10,24,33,34,35,46,47,48,49,56,68,69 |
| | emotions | 24,25,35,43 |
| | recall faces | 23,24,88,101 |
| | recall places | 24,46,49,56 |
| | nouns | 20,50,53,88 |
| 12 | visual test 4 | 1,2,6,10,14,15,19,22,23,24,25,26, 28,33,39,40,41,42,43,44,47,48,51,52,57,60,63,75,79,87,91,92,95,99 |
| | visual test 5 | 2,6,10,14,15,19,22,23,24,25,26 28,33,39,40,41,42,43,44,47,48,51,52,57,60,63,75,79,87,92,95 |
| | visual test 6 | 1,10,41,42,44,47,87,91 |
| | visual test 11 | 1,26,41,87,91 |
| | emotions | 2,10,24,87,91,92 |
| | recall faces | 2,10,22,23,25,26,63,91 |
| | recall places | 10,22,33,41,42,43,45,46,91 |
| | nouns | 10,25,91,92,95 |
| 13 | visual test 4 | 14,15,40,51 |
| | visual test 5 | 17,40 |
| | emotions | 40 |
| | recall faces | 40 |
| | recall places | 40 |
| 14 | visual test 4 | 26,40,62 |
| | visual test 5 | 48,61,62 |
| | visual test 6 | 8,40,48,50,61,62,63,69,79,80 |
| | recall faces | 61,62,63 |
| | nouns | 61,62,63 |
| 15 | visual test 4 | 1,2,3,14,15,19,23,48,49,50 |
| | visual test 5 | 1,3,11,14,15,23,46,50 |
| | recall faces | 3,11,14,15,19,23 |
| | recall places | 11,19,23,41 |
| | nouns | 11,14,15 |
| 16 | emotions | 1,11,19 |
| | nouns | 1 |
| 17 | visual test 5 | 6,8,9,10,13,15,27,28,32,35,39,40,41,42,57,58,59,70,71,72,79,83,87,89 |
| | visual test 11 | 9,13,15,28,32,33,35,39,57,58,59,70,71,72,83,87 |
| | emotions | 6,8,13,24,28,31,32,35,41,42,50,58,71,72,83,87,89 |
| | recall faces | 2,3,6,8,18,24,28,31,32,33,35,39,50,71,72,83,87 |
| | recall places | 59,70,72,83,89 |
| | nouns | 50,51,52,53,89 |

For each patient, electrode leads have numbers uniquely defining the cortical locations.
